# Deep-learning protein structure predictions suggest likely molecular functions for three uncharacterised polytopic membrane proteins from the *P. falciparum* apicoplast

**DOI:** 10.1101/2024.04.13.589297

**Authors:** David L. Murphy, Shahram Mesdaghi, Filomeno Sánchez Rodríguez, Adam J. Simpkin, Luc Elliott, Daniel J. Rigden

## Abstract

Malaria is a burdensome disease to humanity caused chiefly by the still poorly understood parasite genus *Plasmodium*. Much of the pathogenic success of these and other related parasites is due to the presence of the apicoplast, a comparatively poorly characterised biosynthetic organelle containing many proteins of unknown function. Here we present AlphaFold2 protein structure predictions together with further *in silico* analyses to infer molecular functions for the three uncharacterised transmembrane apicoplast proteins PF3D7_0622700, PF3D7_0908100 and PF3D7_1021300. The targets PF3D7_0622700 and PF3D7_0908100 are shown herein to belong to the polytopic Major Facilitator and Cation-Proton Antiporter and Anion Transporter superfamilies respectively, confirming previous suspicions for PF3D7_0908100 of a transporter function. Importantly, our docking screens further suggest pyridoxal-5-phosphate may be transported by PF3D7_0622700, and PF3D7_0908100 likely transports a larger negatively charged metabolite. These findings will help direct experimental assays to confirm what apicoplast metabolites these proteins may transport. PF3D7_1021300 is proposed to possess a six transmembrane alpha-helix domain of a currently unknown fold which may also possess a transporter molecular function. This work highlights the power of high accuracy protein structure predictions to illuminate proteins of unknown structure and function.

## Introduction

AlphaFold2 (AF2) marked the high accuracy ability of deep-learning based methods to predict protein structures [1,2]. Accurate protein structure predictions can aid in the assessment of protein function, and thus confident models derived from next-generation methods have enormous potential to illuminate the molecular function of hypothetical and uncharacterised proteins. Though further efforts have also been made to develop protein language model based methods with state of the art accuracy [3,4], AF2 and previous approaches have relied on the use of residue coevolutionary information present in Multiple Sequence Alignments (MSAs). Such information results from the detection of correlating patterns of covariation between amino acid pairs in MSAs which may be proximal in three-dimensional (3D) protein structure [5], and grants great power in structure prediction. Another advanced bioinformatics method with great potential to aid in the illumination of uncharacterised and hypothetical proteins is HHpred [6,7]. This sequence-based method possesses more powerful abilities to detect distant sequence homology between proteins in comparison to traditionally established sequence searching methods [8]. Much of this homology detecting power results from the use of Hidden Markov Model (HMM) based profile searching aided by (predicted) secondary structure information. We have exploited these plus other recent developments for the molecular function illumination of uncharacterised transmembrane proteins in the apicoplast.

The protozoan phylum Apicomplexa is host to a range of important disease-causing parasites including most notably the malaria causing *Plasmodium* genus. Parasites from the phylum are characterised in part by the presence of the apicoplast, a self-replicating organelle of four membranes known to be a critical biosynthetic powerhouse [9,10].

Alongside a few organellar housekeeping, biogenesis and protein import pathways [11–13], a number of metabolic pathways are known to be present here and are responsible for a small range of required products exported for utilisation elsewhere in the parasite. Various metabolites and inorganic ions are thus required to be imported into the apicoplast to fuel these metabolic activities. The apicoplast like the chloroplast is a plastid, originating from events of cyanobacteria endosymbiosis in an ancient ancestral organism and can vary in their number of membranes, with the four membranes of the apicoplast originating from at least two separate endosymbiotic events [14]. The most popular current theory is that a cyanobacterium (containing two membranes) was endosymbiosed by a primary eukaryote (a red alga) [15,16], which was then endosymbiosed by another secondary eukaryote at which point the last (fourth) membrane was added. Afterwards, the nucleus of the secondary endosymbiont was lost and the photosynthetic function possessed by the organelle was lost through evolution and replaced for parasitism [17]. However, a number of genes have still been retained from the cyanobacterium and alga ancestors, as some apicoplast proteins possess bacterial and algal homologues. It is thought that these protein relics may or may not have been repurposed in function. Only a few genes are encoded within the apicoplast’s chromosome as during evolution most genes were either transferred to the parasite’s nucleus or lost.

Most apicoplast proteins are targeted to the organelle via the secretory endoplasmic reticulum pathway using the bipartite N-terminal targeting sequence region composed of the signal peptide followed immediately by the sequence-variable transit peptide [12,18], and both are ultimately cleaved out [19]. Thanks to a large study by Boucher *et al.* and other previous studies, at least a total number of 346 nuclear-encoded proteins have been experimentally localised to- or predicted to be targeted to the organelle with high accuracy [20]. In terms of the membrane portion of the apicoplast, a number of membrane proteins are known to reside in the apicoplast. Literature surrounding the apicoplast to-date has only described the transmembrane protein type presence of transporters, enzymes and protein import apparatuses, though the presence of currently unknown receptors has also been speculated on for some niche purposes [21,22]. In regards to transporters, a review by Kloehn *et al.* lists all currently known imported and exported apicoplast metabolites of known and unknown transporters [23]. This compiled list totals over 30 known metabolites which lack known transporters. Whilst a number of putative apicoplast transporters from known families exist, it still remains an open question of which metabolites or inorganic ions they may transport. It is also thought that more currently unknown transporters are involved in this puzzle. In conjunction with this, much also remains unknown about the organelle’s composition of inorganic ions.

In another important study, Sayers *et al.* used both experiments and bioinformatics-related data to both suggest (and confirm some of) a list of putative membrane transporters in the *Plasmodium berghei* apicoplast during the blood life stage as candidates for gene essentiality studies [24]. In a guilt-by-association manner a critical number of seven predicted Transmembrane alpha-Helices (TMHs) was used to infer which apicoplast proteins were transporters. The reasoning behind this was based on the fact that a previous group had conducted a bioinformatic analysis of the *Plasmodium falciparum* proteome and found that many of the parasite’s transporters had at least seven TMHs [25,26]. This was in line with the general observation that many transporters from across the biological universe possess approximately eight to fourteen TMHs [27]. The essentiality of these genes during the parasite blood life stage was also factored into account due to the obviously important nature of transporters here in the organelle and their critical role at this life stage.

This research was focused on three selected uncharacterised transmembrane proteins from the Boucher *et al.* high confidence (nuclear-encoded) *P. falciparum* apicoplast proteome predicted to possess multiple (>= 4) TMHs corresponding to PF3D7_0622700, PF3D7_0908100 and PF3D7_1021300 [20]. For these three targets: *i*) PF3D7_0622700 was one of the proteins experimentally localised to the apicoplast by Boucher *et al.*, but was not one of the candidates investigated by Sayers *et al.* most probably due to the possession of less than seven predicted TMHs. *ii*) PF3D7_0908100 and PF3D7_1021300 were experimentally studied by Sayers *et al.* in terms of apicoplast localisation and gene essentiality (mutability) at the blood life stage [24], and amongst the other transmembrane candidates were hypothesised to be transporters. However, in alignment with the results of other large-scale gene essentiality studies in the parasite [28,29], only PF3D7_1021300 has been found to be essential (non-mutable) at the blood stage. All three of these uncharacterised targets currently lack any functional annotation, with no known domains or sequence motifs listed on existing protein databases [30,31]. Potentially existing known structural folds within confident models from deep-learning sources were sought alongside potentially detected distant homology to confirm the transporter functions inferred by Sayers *et al.*. Our logic here was that the three targets could also potentially instead be other polytopic protein types including but not necessarily limited to receptors and proteases, as these can also possess seven or more TMHs, plus could also possess essential or critical functions.

Ultimately, as PF3D7_0622700 and PF3D7_0908100 were found to have copies of the UVB_Sens_prot and Mem_trans domains respectively, we further refer to these proteins as *Pf*_UVBSp and *Pf*_Memtrans respectively. We subsequently next describe background information on these domains and the transporter superfamilies which they fall into. Similarly for PF3D7_1021300, due to our findings suggesting its transmembrane domain is composed of six TMHs, we herein abbreviate this uncharacterised apicoplast protein to UA6TM. Though we had obtained AFDB models of these three targets [32], we also attempted to gain even higher accuracy AF2 models by running AF2 with some slightly different network parameters [33], as discussed shortly in the methods section. However, per-protein only very marginal differences in structure and global pLDDT confidence were observed between the models from these sources. Other AF2 models also obtained in this research primarily correspond to homo-dimer predictions. Slightly prior to the release of AF2, we had also obtained models from other deep-learning based methods such as RosettaFold [34]. However, due to the nature of the results, we only present and discuss findings from a RosettaFold model of UA6TM which augmented the AF2 model centred study. One of the main research end-points for *Pf*_UVBSp and *Pf*_Memtrans as putative transporters were the Small Molecule Docking Screens (SMDSs) we conducted to suggest what native apicoplast metabolites these putative transporters may transport.

The Major Facilitator Superfamily (MFS) transporters constitute a structurally conserved class of secondary transporters enabling the passive facilitated diffusion of organic and inorganic ions across membranes with anion-cation symport, antiport and uniport behaviours [35,36]. These transporters are categorised by having two domains, most commonly within the same chain, interacting via specific TMHs to form an enclosed transmembrane translocation unit. Most MFS domains possess six TMHs meaning that the typical MFS transporter contains twelve TMHs. Upon the update release of Pfam35 in November 2021, the Pfam domain entry (PF04884) for DUF647 was renamed to “UVB_Sens_prot” and categorised into the MFS Pfam clan (cl0015) [37]. In terms of sequence-based relationships with other domains within the clan it possesses none (no HHsearch alignment family pairs of =< 0.1 e-value) [6], and currently lacks an experimental structure. Furthermore, according to the Pfam entry; the domain is only found in eukaryotes and importantly it is most commonly found in protein architectures as the only domain present with one copy. The presence of the domain in this clan would now putatively signify a transporter molecular function, though in all examined literature to-date no molecular function has been hypothesised.

Although the UVB_Sens_prot domain is currently of unknown molecular function, in certain plant-related biological contexts it is known to be involved in maintaining photo-protection and the poorly understood homeostasis of vitamin B6 levels [38–40]. Here, Root UV-B Sensitive (RUS) proteins containing the domain have mostly been documented in *A. thaliana* which possesses six RUS genes (RUS1-6). In all RUS protein related literature analysed to-date the only membrane which is mentioned as the subcellular location of RUS proteins (in *A. thaliana*) has been the chloroplast (envelope) membrane for RUS1, RUS2 and RUS4 [41–43]. Importantly, RUS1 and RUS2 in *A. thaliana* have been demonstrated to physically interact via their corresponding UVB_Sens_prot domains [38,40,44]. Further, the mutation driven disruptions on either gene of these interactions resulted in characteristic phenotypic effects suggesting that their interaction is required for their function.

Vitamin B6 exists in six main forms revolving around the three dephosphorylated (“vitamer”) metabolites corresponding to pyridoxal (PL), pyridoxamine (PM) and pyridoxine (PN); each of which can exist in a mono-phosphorylated form in organisms [45], though pyridoxal-5-phosphate (PLP) is the main biologically important form of the vitamin required as a cofactor for biosynthesis enzymes such as Aminotransferases (ATs). Only a very few transporters of the vitamin are known in eukaryotes and very little literature exists on its transport in organisms [46]. It is known in plant organisms such as *Arabidopsis thaliana* the only currently known vitamin B6 transporter is the PUP1 transporter of the three vitamers which is currently only known to reside in the cell membrane [47]. Many of the experimental observations would positively support RUS proteins as transporters of vitamin B6 molecules, though the two most critical are: *i*) An exogenous additional supply of all of the vitamers is able to at least partially recover the RUS mutant phenotypes [40]. *ii*) That RUS mutants have decreased cellular levels of all six forms of the vitamin. Recently Tong *et al.* determined additionally that both RUS1 and RUS2 physically interact with ATs in *A. thaliana* [44]. The authors speculated that though in the current understanding of the vitamin’s homeostasis the binding of PLP to proteins such as ATs is thought to sequester PLP, the RUS-AT binding may also drive conformational changes in the ATs’ binding sites affecting the ability of PLP to bind to the enzyme, which in turn would affect levels of the metabolite. Though it is not our prerogative to argue centred on evidence that UVB_Sens_prot is a transporter of vitamin B6 metabolites; we simply reason here based on the observations that the homeostatic mechanism by which RUS proteins work is instead more likely resultant from them acting as secondary transporters of the vitamin in organisms. We must clarify here that in all of the relevant literature we have consulted to-date, essentially no publications have discussed the fact that the UVB_Sens_prot domain belongs to the MFS and that this renders the RUS protein family a putative group of transporters.

One critical apicoplast metabolite which was not listed by Kloehn *et al.* for unknown reasons is PLP. The apicoplast possesses metabolic pathways to synthesise the Iron-Sulphur clusters required internally as prosthetic groups [48,49], and PLP is required as a cofactor for this in the *P. falciparum* apicoplast [50]. It is known that *P. falciparum* can biosynthesise PLP via the two two routes of either: *i*) *de novo* biosynthesis from glutamine and a phosphate donor substrate via the enzyme complex of Pyridoxine biosynthesis proteins (Pdx1 and Pdx2) [51,52]. Or *ii*), dephosphorylated vitamer salvage from host followed by phosphorylation-modification by the enzyme Pyridoxal Kinase (PdxK) [53]. All three of these enzymes were only found to be localised to the cytosol in the parasite [54]. Only a single (non-recent) review was identified from literature searches regarding what vitamin B6 transporters may exist in the parasite and it purely describes the fact that there is no known PLP transporter in the *P. falciparum* genome [55]. Thus, it would appear that the apicoplast requires transporters in order to receive PLP and we later dissect the plausibility of *Pf*_UVB_Sens_prot as a transporter of the vitamin.

The Mem_trans domain (Pfam PF03547) belongs to the Cation Proton Antiporter and Anion Transporter superfamily (Pfam clan “CPA_AT”, cl0064), which we herein abbreviate to CPAS. The superfamily is diverse in terms of the sequence [56], structure and the range of substrates transported by its members. The CPAS has traditionally been associated with the transport of both organoacids and inorganic ions (predominantly sodium and hydrogen ions) in an antiport or symport manner of secondary transport [57]. The structural diversity within the CPAS is predominantly resultant from the variation in topology, the number of TMHs, and the heterogeneity of functionally important alpha-helices within the transmembrane regions of the domains. However, one consistency between the CPAS families forming a superfold is that the domains contain a partitioning into two subdomains termed the scaffold and core regions typically formed by the intertwining of two internal repeats of alpha-helices corresponding to the N- and C-terminal halves. The approximate interior structural gap caused by this partitioning produces the transmembrane translocation (transport) region [58,59], a type of behaviour which has also been observed in other membrane transporter superfamilies [60]. In this translocation region of CPAS transporters, non-canonical forms of TMHs exist such as broken alpha-helices (BHs) and re-entrant alpha-helices [61], as has been found in other types of membrane transporters [62], and are important for ligand binding and transport.

The Mem_trans family is more commonly referred to as the “Auxin Efflux Carrier” (AEC) family and is listed on Pfam as existing in a wide range of eukaryotic and bacterial phyla. It is characterised mostly from plant organisms where the transport of auxin is known [63]. To-date, we have only observed in plant-related literature subcellular localisations of the endoplasmic reticulum and cell membranes [64]. However, members of the AEC family have been found in cyanobacteria [65]. In other bacterial contexts, sub-groups of the family are also associated with the transport of a few other certain small organic substrates. More specifically, the MleP transporters are associated with malate and malonate [66,67], and CitP transporters with citrate [68]. Proton antiport has traditionally been thought to occur in the family, though it has only been experimentally confirmed in the family for CitP transporters [69], and in the recent three structure solving studies of the PIN auxin transporters a lack of signs of the co-transport is described but not necessarily ruled out [70–72]. At the time of conducting this research no experimental structures of the Mem_trans domain existed, although a topology had been proposed in 2021 by Sudha *et al.* based on a trROSETTA structure prediction study of the CPAS [61,73]. However, very recently three publications covering a total of nine cryo-electron microscopy structures of the AEC family have been released in the Protein Data Bank (PDB) [70–72,74].

## Methods

### Data resources

Protein fasta sequences were downloaded from PlasmoDB (release 39) [30]. The HHsuite compatible PDB70 [74], PfamA (release 30/01/2019) [37], and UniProt20 (release 20/02/2016) were downloaded for local use with the suite [31]. Pfam domain seed MSAs were downloaded from the 35th version of the online database [37]. The RCSB mirror of the entire PDB database of protein structures was downloaded on 28/08/2020 plus then formatted by the DALIitev5.0 suite scripts import.pl and format.pl respectively for DALI utilisation [74–76]. This same local PDB database was also formatted for GESAMT utilisation via the program itself with the “--make-archive” option [77]. All PDB structure codes from the weekly version release 21/08/2020 of the Transmembrane PDB (PDBTM) database were used for the structure searches [78].

### Sequence-related bioinformatics

Sequence database similarity searches were conducted by: *i*) HHblits against the (HHpred compatible) UniProt2016_20 sequence database [8,31,79], using a search with eight iterations and e-value cut-off of 0.001 (all other parameters kept at default). *ii*) PSI-BLAST against the Non-redundant (NR) sequence database via the NCBI BLAST webserver [80,81], with three search iterations and an e-value threshold of 0.001 (all other parameters kept at default). *iii*) HHsearch against the above-mentioned PfamA domain and PDB70 structure databases in the default manner (e-value cut-off 0.001) using the HHblits generated MSAs from *i*) [6,37,74]. *iv*) HHpred against the *Toxoplasma gondii* (TGME49), *A. thaliana* (TAIR) and *P. falciparum* (3D7) proteomes (via the MPI toolkit webserver) in the default manner [7,82]. Jalview (version 2.11.1.3) was used for the visualisation of sequence alignments [83]. Sequence-based TMH predictions were conducted by the TOPCONS2 webserver [84].

### Transit peptide boundary identification and removal

The sequence position of transit peptide boundaries were estimated by analysing towards the N-termini: *i*) Approximately where higher sequence concentrations of lysine and arginine residues were. *ii*) Where AFDB model regions were more distant from the rest of the chain’s structure and possessed unfolded structure. These signal and (estimated) transit peptide removed versions of the protein sequences were then used for the AF2 structure prediction protocol described below, plus these same regions were removed from the AFDB models as truncated versions. Thus all AF2 models used correspond to the sequence regions in *Pf*_UVBSp, *Pf*_Memtrans and UA6TM of 68-381, 86-476 and 69-454 respectively. For the RosettaFold structure prediction of UA6TM conducted earlier in our research, residues 82-454 were modelled in accounting for a slightly different estimated transit peptide boundary based on other previous bioinformatics analyses not described further.

### Structure prediction

RosettaFold structure prediction of UA6TM was conducted by the Bakerlab webserver [34]. AlphaFold Database (AFDB) release version 1.0 models were downloaded from the official AFDB web-portal [1,32]. Monomeric AF2 structure prediction was conducted using a non-Docker implementation of the official AlphaFold version 2.0.0 code-base and required databases [85]. Five models were generated using the default CASP14 network options and parameters for the sequence database search, structure prediction and AMBER refinement steps except for: *i*) Five sequence database search iterations were conducted by both hhblits and jackhmmer from a default of one and three respectively. This was achieved by appropriately editing lines 39 and 38 in the AF2 code-base “alphafold/data/tools/” directory scripts hhblits.py and jackhmmer.py respectively. *ii*) A number of 24 recycles was used by modifying the “num_recycle” value of the model config dictionary in the AF2 code-base script “alphafold/model/config.py” at lines 106 and 399 from the default of three. The AF2 advanced (google) colab webserver tool (https://colab.research.google.com/github/sokrypton/ColabFold/blob/main/beta/AlphaFold2_advanced.ipynb) was used to generate homo-dimer models [86]. The options “use_turbo” and “use_ptm” (pTM network) were unselected [33], mmseqs was used for the MSA generation step [87,88], with all other options and parameters the default. Additional modified runs were also conducted where the structure sampling parameter “num_samples’’ was increased from 1 to 16. The ColabFold AF2 (google) colab webserver tool v1.3.0 was used (for monomeric structure prediction of UA6TM) in the default manner.

### Structure alignment/searching

GESAMT (v1.5) from the CCP4 suite (v7.0) and the script dali.pl (DALI) from the DALIlitev5.0 suite were used to conduct structure similarity (structure alignment) searches against the PDBTM [76–78,89]. The DALI hierarchical search mode (“--hierarchical” option) was used, and the GESAMT “-high” option was used for (marginally) higher structure search selectivity and sensitivity, but all other GESAMT parameters and options were the default. The FoldSeek webserver was used to conduct structure searches against the entire AFDB database (third release) of structure predictions by both the 3Di and TMalign modes [90,91]. The DALI webserver was used to search UA6TM models against the entire PDB on March 18th 2024 [92]. TMalign was used in the default manner to conduct structure alignments for fold comparison [91].

### Structure analyses

AF2 models were coloured in terms of confidence (pLDDT) by the “colour_af_plddt.py” PyMol script according to the official AFDB per-residue pLDDT (confidence) scheme [32,93]. Conservation was principally determined and expressed on models by use of the (previous-generation) CONSURF webserver in the default manner by use of the AF2 generated MSAs alongside the top models as input [94]. Charges of model surfaces were determined by the use of the APBS electrostatics plugin of Pymol (version 2.4.2) in the default manner [95], with the protein molecule prepared by pdb2pqr method and a charge range of +/- 5.00 [96].

### Small molecule docking screens

Models were prepared for docking by adding hydrogen atoms and charge using the Chimera (UCSF) DockPrep tool in the default manner [97]. The side-chain terminal atom coordinates of specific residues from the later discussed analyses were used as grid centres. The Webina webserver (v1.0.3) 3D protein structure visualiser tool was used to aid in the process of the selection of the grid dimensions (x, y, z) in size of Ångstroms to cover the transport regions of interest in the prepared model [98]. The grid determined for the target protein via visualisation in Webina was the same grid specified for the target protein SMDS conducted via the SWISSDOCK webserver [99,100]. A relatively larger 3D docking grid was designated for both SMDSs in order to cover all of the transmembrane translocation regions’ interior surfaces. We list this docking grid information used in table 1.

**Table 1:** Docking grid information.

| **SMDs** | **Residue** | **Structural location** | **Side chain atom (grid centre)** | **Dimension of grid (x, y, z) Å** |
| --- | --- | --- | --- | --- |
| *Pf_UVBSp* | ARG143 | TMH3 (first chain),<br>at back of pocket | NH2 (nitrogen) | 42, 50, 42 |
| *Pf_Memtrans* | HIS247 | TMH4 (first BH),<br>tunnel region | ND1 (nitrogen) | 25, 40, 25 |

The molecule versions of the metabolites present at physiological conditions were identified on the PubChem database (displayed in table 2) [101], and their 3D conformational structure files in SDF format were downloaded from the corresponding entry pages [102]. These molecule SDF structure files were used for Webina docking. For SWISSDOCK these 3D SDF structure files of the molecules were converted in Chimera to MOL2 format 3D structure files, but were checked after the conversion to ensure that they possessed the same atoms and bonds (including for order). As the Coenzyme-A and De-phospho-Coenzyme-A Pubchem database entries did not have 3D conformer versions available, the two-dimensional versions of the molecule structures were downloaded in SDF format instead. The online openbabel incorporating chemoinformatics conversion tool (http://www.cheminfo.org/Chemistry/Cheminformatics/FormatConverter/index.html) was then used to generate the full 3D coordinate files in SDF format [103]. In Webina, the recommended (default) high exhaustiveness value of the docking search of 8 was used and similarly in SWISSDOCK the accurate mode was used with no (zero Å) flexibility allowed for the protein residue side chains.

**Table 2:** List of metabolite molecule versions used in SMDSs.

| Metabolite | Abbreviation | Docked to | Molecular weight (g/mol) | Formal charge | Molecular complexity | Pubchem compound ID |
| --- | --- | --- | --- | --- | --- | --- |
| Pyridoxal | PL | *Pf*_UVBSp | 167.16 | 0 | 162 | 1050 |
| Pyridoxamine | PM | *Pf*_UVBSp | 169.20 | +1 | 143 | 25245492 |
| Pyridoxine | PN | *Pf*_UVBSp | 169.18 | 0 | 142 | 1054 |
| Pyridoxal 5'-Phosphate | PLP | *Pf*_UVBSp and *Pf*_Memtrans | 245.13 | -2 | 281 | 644168 |
| 5-aminolevulinic acid | 5-ALA | *Pf*_Memtrans | 131.13 | 0 | 116 | 7048523 |
| Adenosine Di-Phosphate | ADP | *Pf*_Memtrans | 424.18 | -3 | 618 | 7058055 |
| alpha-Keto-Glutarate | aKG | *Pf*_Memtrans | 144.08 | -2 | 160 | 164533 |
| Aspartate | ASP | *Pf*_Memtrans | 132.09 | -1 | 122 | 5460294 |
| Adenosine Tri-Phosphate | ATP | *Pf*_Memtrans | 503.15 | -4 | 772 | 5461108 |
| Biotin | (None) | *Pf*_Memtrans | 243.31 | -1 | 292 | 171548 |
| Citrate | (None) | *Pf*_Memtrans | 189.10 | -3 | 211 | 31348 |
| Coenzyme-A | CoA | *Pf*_Memtrans | 763.5 | -4 | 1240 | 25113190 |
| De-phospho Coenzyme-A | DeP-CoA | *Pf*_Memtrans | 685.5 | -2 | 1080 | 45266562 |
| Dimethylallyl Pyrophosphate | DMAPP | *Pf*_Memtrans | 243.07 | -3 | 279 | 15983958 |
| deoxy Thymidine 5'-Triphosphate | dTTP | *Pf*_Memtrans | 478.14 | -4 | 825 | 11988283 |
| Glutamate | GLU | *Pf*_Memtrans | 146.12 | -1 | 134 | 5460299 |
| Thiamine Pyrophosphate | TPP | *Pf*_Memtrans | 422.29 | -2 | 548 | 15938963 |
| Nicotinamide Adenine Dinucleotide | NAD+ | *Pf*_Memtrans | 662.4 | -1 | 1110 | 15938971 |
| Nicotinamide Adenine Dinucleotide Phosphate | NADP+ | *Pf*_Memtrans | 740.4 | -3 | 1270 | 15938972 |

Hydrogen atoms were re-added to the output Webina ligand poses in Chimera, where they were then exported with the prepared protein model structures to individual protein-ligand complexes. Coordinate data of the output SWISSDOCK ligand poses was individually appended to the prepared protein model structures by scripts to form the complexes. The CSM-lig and Prodigy-Ligand webservers were also used to obtain more accurately Predicted Binding Affinities (PBAs) for all of these individual protein-ligand complex structures [104,105]. We refer to this as rescoring. Canonical SMILES strings from the corresponding metabolite PubChem entry pages were used for CSM-lig rescoring [106]. Prodigy-Ligand PBA energies in terms of the free energy of ligand binding (ΔG) are equivalent to those PBA energies from Webina and SWISSDOCK in kcal/mol; whereas, the output of CSM-lig includes the PBA in relative terms of the binding association constant -log_10_(K_d_/K_i_). With respect to the latter, more positive values indicate higher PBAs. In our analyses we have focused only on (and rescored) the ligand poses which had bound in the transmembrane translocation regions, and even more specifically for *Pf*_Memtrans-only the ligand poses which had bound in the tunnel region. The standalone LigPlot (version 2.2.5) tool was used to determine the ligand-residue interactions of the individual protein-ligand complexes [107].

## Results and discussion

### Pf_UVBSp

#### *Pf*_UVBSp adopts to the classic MFS transporter structure

AF2 models of *Pf*_UVB_Sens_prot (figure 1a) possessed a transmembrane domain of six TMHs which in terms of surface possessed a slightly concave shape marked by a prominent central pocket at one side to the bundle (depicted shortly). However, as importantly also represented in figure 1b, an intrinsically disordered Large Loop Region (LLR) exists between the fourth and fifth TMHs at residues 219 to 324 with very low pLDDT. The AFDB model possessed the higher average model pLDDT confidence score (including with the LLR removed), and possessed a confident transmembrane region pLDDT score of 73.74. This model was thus used for all *Pf*_UVB_Sens_prot model-based analyses except for CONSURF-derived conservation and the vitamin B6 SMDS.

**Figure 1:**
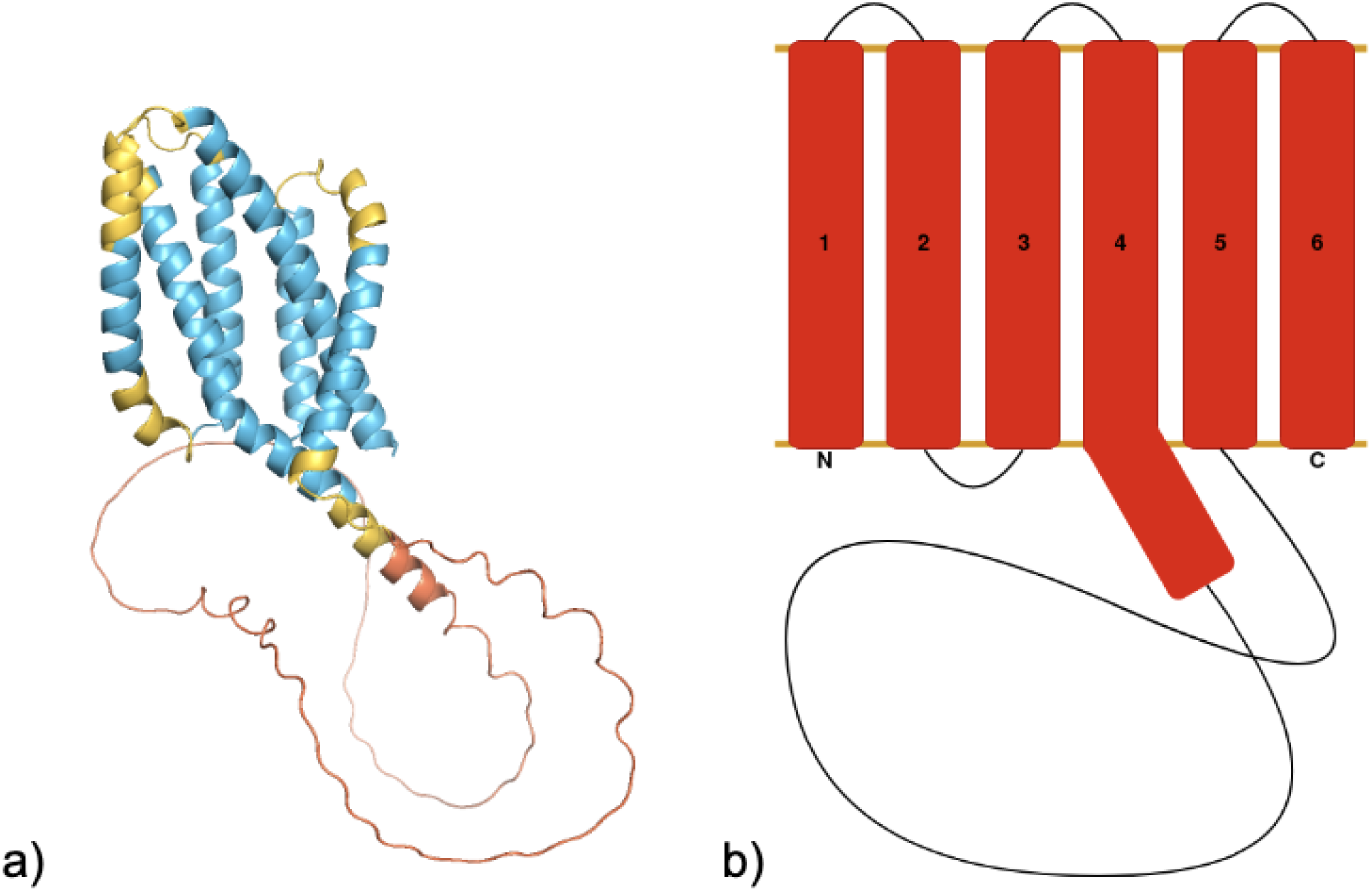
Predicted *Pf*_UVBSp structure and topology. a) *Pf*_UVBSp AFDB model coloured by the official AFDB (per-residue) pLDDT confidence scheme [32]: Light red (very low pLDDT < 50), yellow (low pLDDT 70 > 50), light blue (high/confident pLDDT 90 > 70), deep-blue (very high pLDDT > 90). b) Cartoon of *Pf*_UVBSp membrane topology: TMHs not drawn to their 3D location in model structure, alpha-helices red, loops black and membrane bi-layer boundaries gold, N- and C-termini are also depicted.

The DALI and GESAMT structure searches of these AF2 models against the PDBTM revealed only relatively high-scoring structure hits from the MFS and no other transmembrane families [76–78]. Here, the highest DALI Z-scores and GESAMT Q-scores achieved by these hits were 11.8 and 0.14 respectively, and in table 3 we have listed the highest Z-score hits representative of the MFS (Pfam) family they belong to [37]. As demonstrated in figure 2a, the MFS hit alignments covered the whole of the model transmembrane regions with significant similarity (including identical topology). Yet, as also seen in figure 2b, only half of the transmembrane region of the MFS structures was covered corresponding to only one of the two domains forming the MFS transporters. The use of TMalign to align the AFDB model to the second (MFS) domain of the chains of a few of these higher scoring MFS structure hits revealed TMscores of around 0.6 and 0.7 [91], even with the LLR kept in the model, definitively signifying a fold match. It is thought based on the LLR in these models plus possession of only one MFS transporter domain that these hits would otherwise have achieved much higher Z- and Q-scores for a transmembrane protein of this size and the high similarity observed in superimpositions.

**Figure 2:**
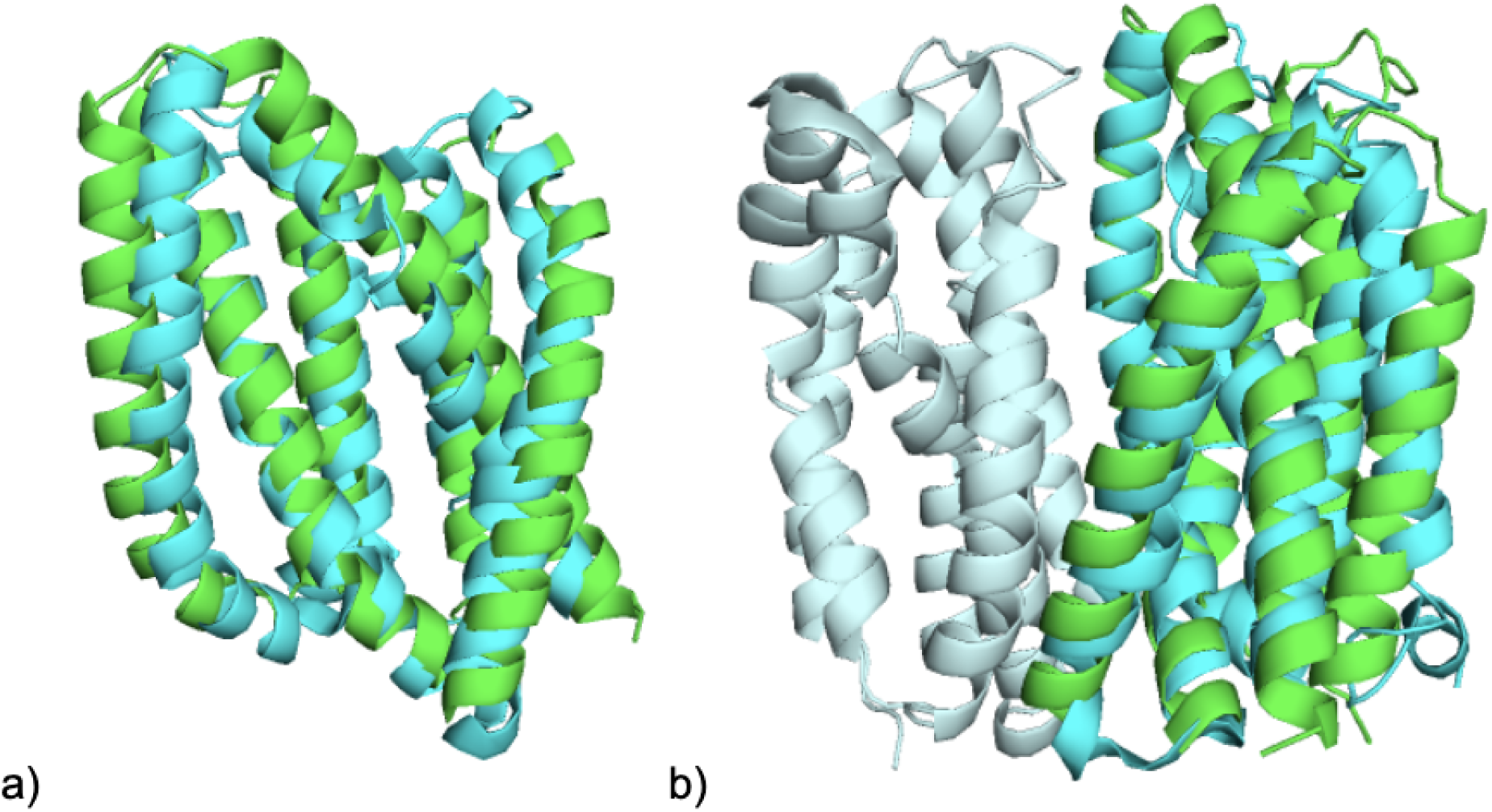
MFS transporter structure superimposition of *Pf*_UVBSp a) *Pf*_UVBSp truncated AFDB model (with LLR and end helical extension of TMH4 omitted) in green superimposed from DALI alignment on the second (MFS) LacY_symp domain of 4oaa-A in cyan. b) The same superimposition but showing the entire LacY transporter structure with the first and second domains of 4oaa-A coloured pale cyan and cyan respectively [108].

**Table 3:** DALI identified top scoring structure representatives of MFS families.

| Pfam domain | DALI hit (protein name) | Chain ID | DALI Z-score |
| --- | --- | --- | --- |
| LacY_symp | Lactose permease | 4oaa-A | 11.8 |
| FPN1 | Ferreportin | 5aym-A | 11.2 |
| MFS_1 | L-fucose-proton symporter | 3o7q-A | 11.1 |
| Sugar_tr | Solute carrier family 2, facilitated glucose transporter member 5 | 4yb9-D | 8.7 |

Homo-dimer models of *Pf*_UVB_Sens_prot were further obtained by the AF2 advanced colab webserver to confirm that the MFS structure possessed by this target could form the full transporter unit [86]. Indeed, at least for the top ranked homo-dimer model the MFS structure was very well recapitulated and it was observed that both monomers bind at the correct two TMHs (the second and fifth TMHs), plus with the same approximate positioning as how the domains in MFS transporters bind to each other. This was confirmed by the structural alignment-superimposition of the model on structures of these transporters. GESAMT was required to simultaneously structurally search the two chains of the homo-dimer model (with its LLR removed) against the PDBTM and it was found to have high global structural similarity to MFS transporter structures. The highest scoring hit (of Q-score 0.32) here corresponded to the *E. coli* lactose permease (LacY) structure 5gxb-A [108], and the homo-dimer model superimposition here is displayed in figure 3. Thus, these homo-dimer models suggest that *Pf*_UVBSp homo-dimerisation to form an MFS transporter is feasible.

**Figure 3:**
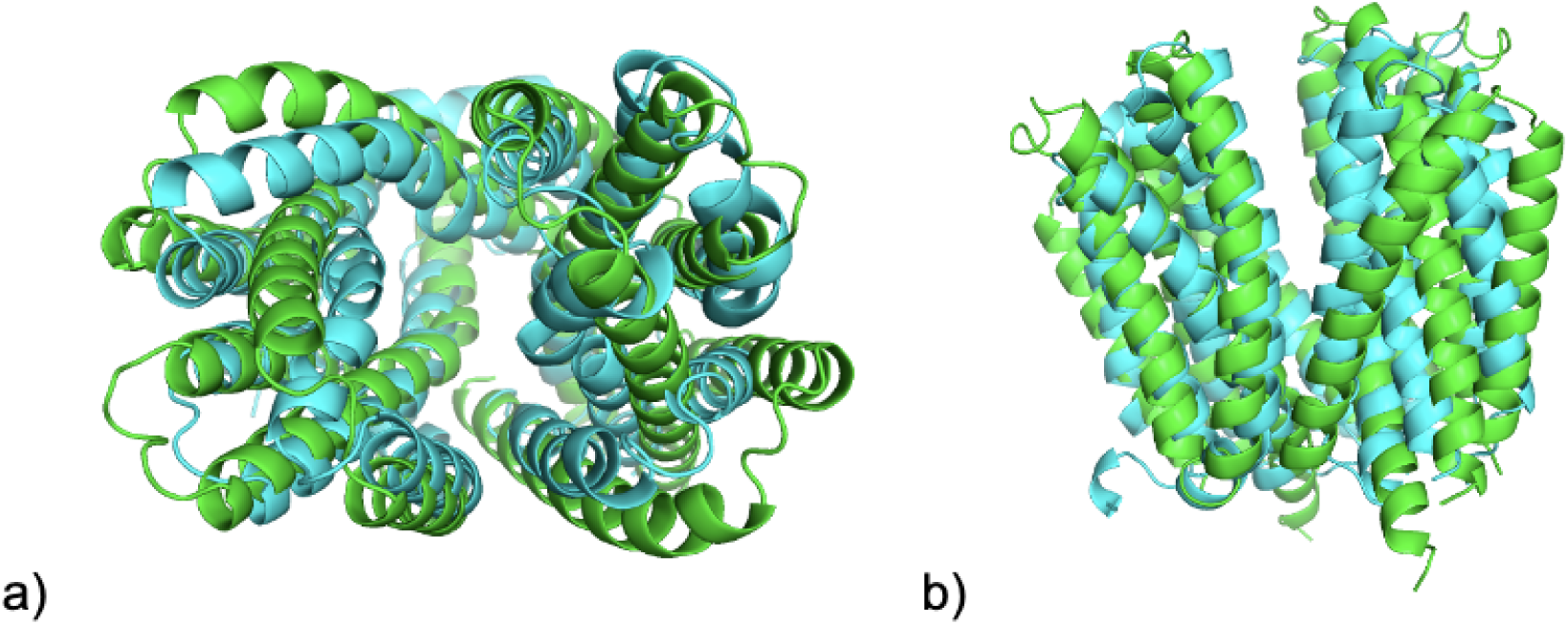
MFS transporter structure superimposition of *Pf*_UVBSp homo-dimer. Superimposition of *Pf*_UVBSp AF2 homo-dimer top model (green) with 5gxb-A (cyan) from GESAMT alignment. View from a) top of TMH bundle, and b) from side.

#### *Pf*_UVBSp contains a single copy of the UVB_Sens_prot domain

In the *Pf*_UVBSp HHsearch sequence search against the Pfam domain database [6,37], UVB_Sens_prot was the only homologous domain hit as marked by a significant HHpred probability score of greater than 70 (table 4), and no sequence similarity was found with known structures from the tool’s PDB70 search [74]. Approximately the first third of the *Pf*_UVBSp sequence after the approximate transit peptide region was aligned to the first half of the UVB_Sens_prot domain representative protein CRAGI’s (UniProt ID: K1RDB8) sequence. When this aligned region of the sequence is displayed on the AFDB model it corresponds to the first four TMHs, up to immediately prior to the LLR. However, as also displayed in table 4, a repeat search here with the LLR removed from the sequence resulted in a significant improvement in the representative hit’s e-value, and instead covered virtually all of the transmembrane *Pf*_UVBSp and CRAGI sequence regions. However, as originally found with the whole sequence version, HHblits-Uniprot and PSI-BLAST-NR sequence searches with the LLR sequence region removed still did not identify homologous sequences outside of *Plasmodium* [31,79–81], meaning that the domain’s lack of typical sequence-based detection is not purely due to the LLR. HHpred sequence searches of the same LLR-removed sequence against the *T. gondii* proteome also did not reveal any distant homology to other proteins which may contain the domain [7,82]. Though such a finding may somewhat contribute in hinting towards a lack of the domain’s existence in other Apicomplexans, we have not effectively investigated such a possibility further.

**Table 4:** *Pf*_UVBSp HHsearch/HHpred hits with UVB_Sens_prot-containing proteins.

| Hit | Probability score | e-value | Percent identity (%) | Similarity score | Query residues | Target HMM residues | Target HMM size |
| --- | --- | --- | --- | --- | --- | --- | --- |
| CRAGI | 95.59 | 0.022 | 13 | 0.181 | 69-198 | 26-154 | 241 |
| *CRAGI | 98.17 | 2.4e-05 | 15 | 0.215 | 69-379 | 26-220 | 241 |
| *RUS4 | 98.09 | 2.6-e04 | 17 | 0.267 | 58-379 | 117-328 | 520 |

\*Alignment to hit excludes residues 219-324 removed in accounting for the LLR.

Contrastingly, the HHpred search of the LLR removed sequence against the *A. thaliana* proteome revealed two protein sequences to have e-values of less than 0.001, which corresponded to RUS3 and RUS4. The highest scoring and most sequence-similar of these hits was RUS4 (UniProt ID: Q67YT8), which is also displayed above in table 4. This alignment showed a higher number of similar and identical residue positions in comparison to that of the domain representative CRAGI, and these residues in *Pf*_UVBSp which were within the pocket plus the interior region surrounding it are shown in figure 4. Many of these residues were structurally equivalent or at very similar TMH positions in both protein AFDB models. In line with later discussions, this suggests that a similar ligand may bind to both. In any regard, it would appear that *Pf*_UVBSp possesses plant homologues within the RUS family, however this is disputed next.

**Figure 4:**
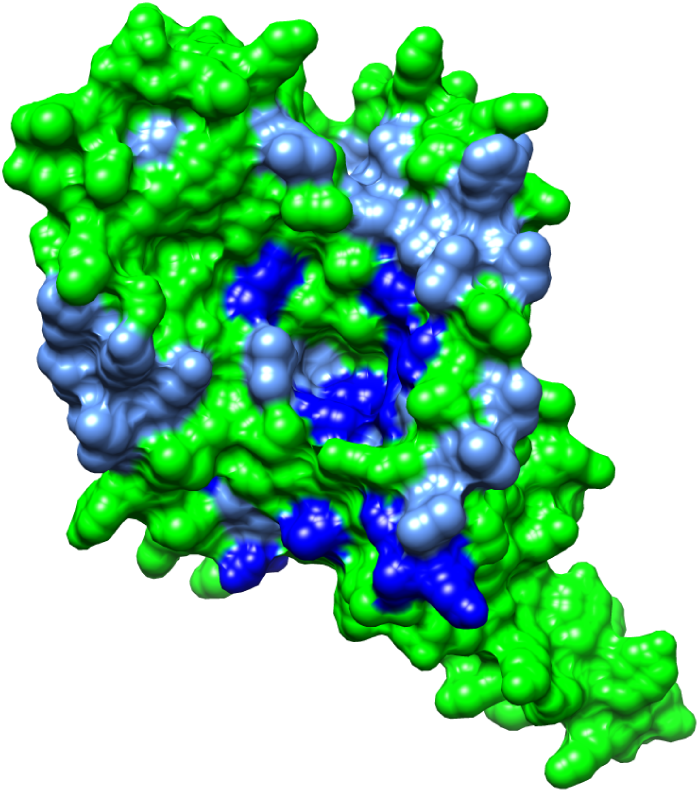
RUS4 sequence similarity within *Pf*_UVBSp pocket. The HHpred-derived pairwise alignment identical (deep-blue) and similar (light-blue) residues between the *Pf*_UVBSp and RUS4 sequences expressed on the surface of the (LLR removed) *Pf*_UVBSp AFDB model in green.

Importantly, it must be highlighted here that regardless of their corresponding organisms many RUS proteins are currently annotated on UniProt with UVB_Sens_prot being the only domain present [31]. However, it has been observed in quite a number of their AFDB models that the C-terminal regions outside of the transmembrane domain contain an alpha-beta domain; which differ reasonably in terms of structure between the RUS proteins- to the extent that they cannot by visual means be deemed as adhering to certain types of alpha-beta architecture. For the above-mentioned RUS4 we have extracted the sequence of this region (residues 328-520) and used HHsearch to search against Pfam with, for which no distant homology to known domains was found. Due to the lack of mention in RUS family-related literature, we are unsure of what the significance of these unknown domains are in terms of protein function, though we have not investigated them further. Hypothetically, the previously discussed interactions between RUS and AT proteins in *A. thaliana* could even be mediated by these uncharacterised domains [44]. It is still possible that due to the plant chloroplast and apicoplast both possessing cyanobacterial endosymbiotic evolutionary ancestry these two different protein families containing the UVB_Sens_prot domain could have originated from a distant common ancestor. Subsequent events in evolution could have caused loss of this C-terminal domain from these apicoplast proteins in *Plasmodium*.

Due to the presence of only a single copy of the UVB_Sens_prot domain, to form a full MFS transporter *Pf*_UVBSp must either homo-dimerise or hetero-dimerise with the UVB_Sens_prot domain of another protein, potentially as an instance of homologues analogous to the RUS1 and RUS2 proteins in *A. thaliana* [38,40,44]. Though the previously discussed sequence database searches did not suggest the presence of another *Pf*_UVBSp homologue within the *P. falciparum* genome, HHpred was anyhow used to search the LLR-removed sequence version against the *P. falciparum* proteome. It was found in the output of hits that besides the *Pf*_UVBSp self-sequence match, only a very few local portions of MFS transporters containing different MFS domains with very poor probability and e-value scores were identified. In alignment with the previously discussed confident homo-dimer model(s) adhering to the MFS transporter structure, this leads us to contend that unlike the hetero-dimerisation scenario of the RUS1 and RUS2, *Pf*_UVBSp homo-dimerises to fulfil the formation of a MFS transporter.

#### Docking confirms vitamin B6 as a plausible *Pf*_UVBSp transport substrate

As displayed in figure 5c, the use of the PyMoL APBS electrostatics tool revealed the *Pf*_UVBSp AF2 models to possess interior surfaces which are mostly more positively charged [95], including where the putative ligand-binding pocket is. Also shown in figure 5a, the use of the CONSURF conservation calculation tool additionally revealed the transmembrane domain to be highly conserved [94], including in terms of the residues forming the interior region’s surface (figure 5b). Many of the most strongly conserved residues in this region were positively charged, suggesting that the positive charge in and around the pocket’s surface may be critical for ligand binding. Some of the key conserved residues are discussed shortly in the context of our SMDS.

**Figure 5:**
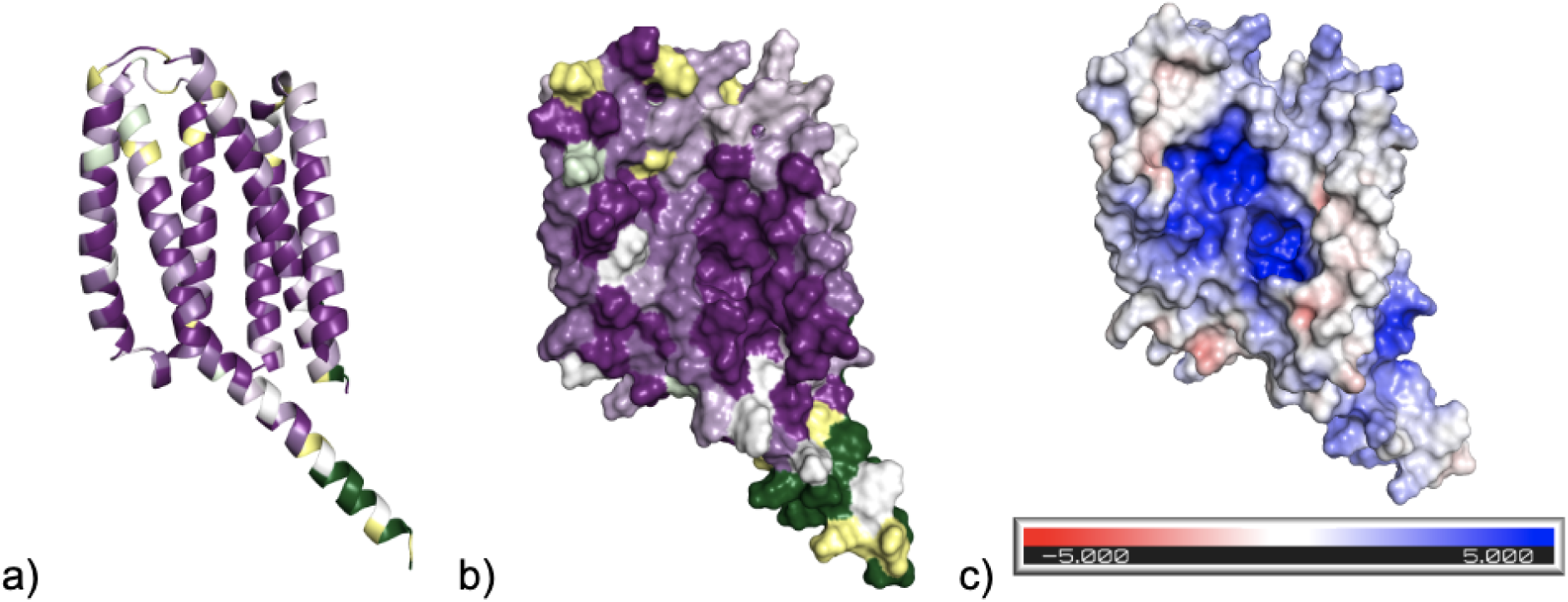
*Pf*_UVBSp conservation and surface electrostatics. a) Cartoon and b) surface representations of AF2 top model coloured by the CONSURF conservation scheme [94]: Residues are coloured yellow where no MSA depth (gaps) exists and a conservation score has not been calculated. All other residues ranging in the colours from dark green to white to dark purple correspond to residues ranging in conservation scores from variable (1) to strongly conserved (9) respectively. c) PyMoL APBS electrostatics surface depiction with charge scale bar of the AFDB model (LLR removed) [95,96].

Due to the previously described connection of plant RUS proteins to vitamin B6 and the presence of multiple equivalent alignment-identical residues to RUS4 in the pocket region, PLP plus the three vitamers (table 2) were specifically docked to the *Pf*_UVB_Sens_prot homo-dimer top model [98–100]. We note here that for unknown reasons CSM-lig predicted PBAs of a large proportion of the *Pf*_UVBSp vitamin B6 metabolites ligand poses with suspicious negative values [104]. However, all had otherwise attained negative PBA energies of lower than -3.0 kcal/mol (including from Prodigy-Ligand rescoring) [105], and in our analyses we have ignored all negative CSM-lig PBAs. As displayed in figure 6, docked poses of all four vitamin B6 metabolites in the transmembrane translocation region presented viable negative PBA energies of around -5 to -8 kcal/mol. The corresponding range of (non-negative) CSM-lig PBAs were of between 25.5 and 67.7. However, nearly all ligand poses with higher CSM-lig PBAs of greater than 50 corresponded to SWISSDOCK ligand poses of PLP and PL. Whilst more negative SWISSDOCK PBA energies were attained for PLP poses, relatively similar PBA energies were achieved between all forms of the vitamin.

**Figure 6:**
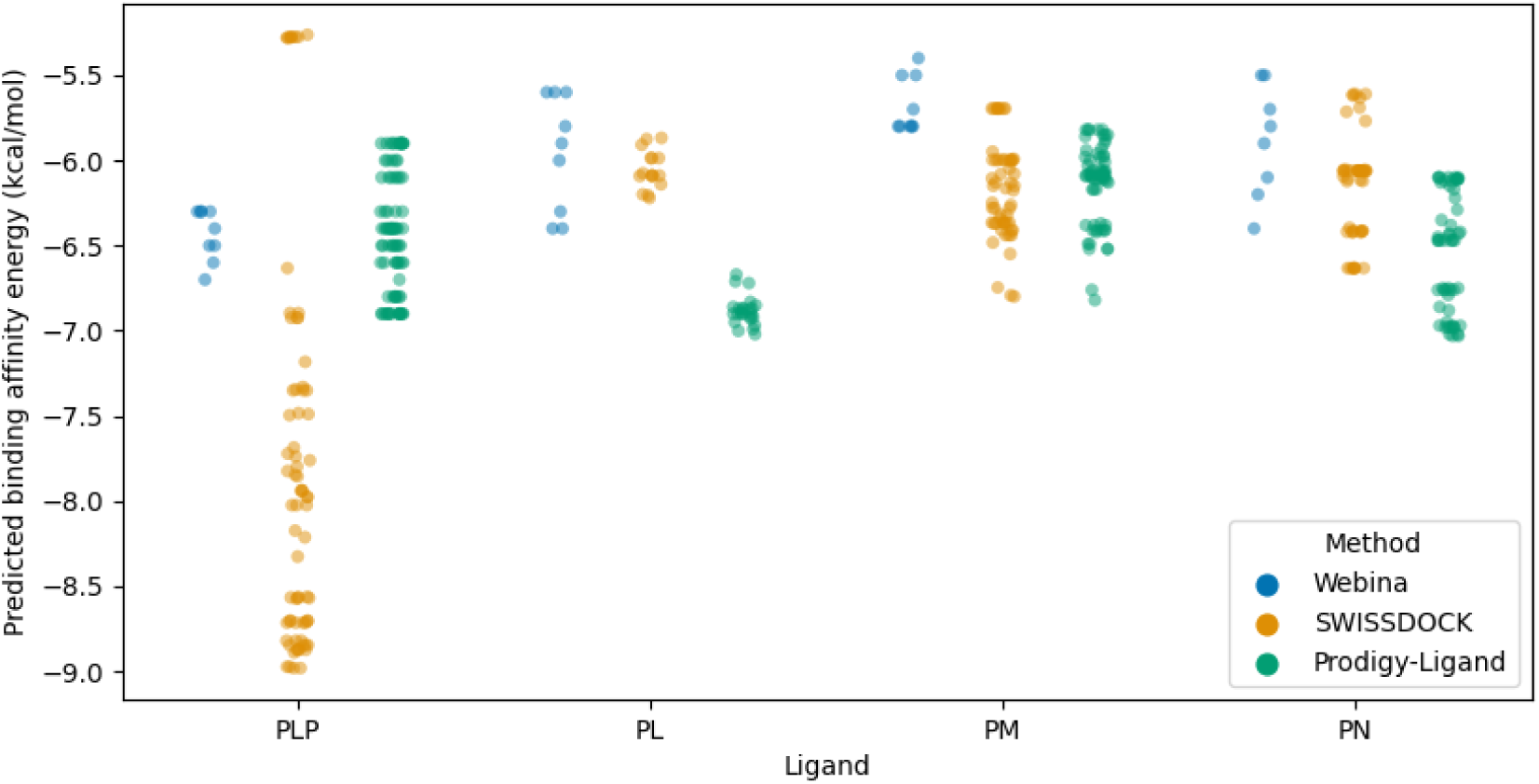
PBA energies of *Pf*_UVBSp vitamin B6 SMDS. Webina, SWISSDOCK and Prodigy-ligand PBA energies of all four vitamin B6 metabolites docking poses which had bound within the *Pf*_UVBSp transmembrane transport region.

Importantly, for the ligand poses from both docking methods there was little difference in the PBAs from all four sources of PBA regardless of whether the ligand had bound in the pocket regions or elsewhere in the transmembrane translocation region. Another observation in line with this was that there were no discernable trends in the PBAs of the vitamin B6 metabolites in terms of the ligand poses binding modes in the pocket region (i.e., no particular orientations of the poses were found to have better PBAs than the others), though we next further discuss these pocket region binding modes.

All of the vitamin B6 metabolite ligand poses from Webina which had bound in the pocket regions bound in two approximate modes where the ligand’s pyridine ring was oriented with regards to the position of the phosphate group (not present in the vitamers) facing either the back of the pocket (inwards) as in figure 7, or outwards from the pocket. Such a form of binding was interesting as it is highly similar to how ligands in MFS transporter experimental structures are approximately found to bind to their corresponding pockets. However, only a few of the SWISSDOCK PLP, PL and PN ligand poses which had bound in the pocket region were similar to these two approximate binding modes. A particular trend here found by the use of the LigPlot tool was the hydrogen bonding of the ammonium groups on the arginine residue side chains to the oxygen atoms on the phosphate group of PLP. This was most notable for the residue ARG143 at the back of the pocket for this ligand’s inwards oriented poses. In equivalent ligand poses where the ligand’s pyridine ring group was oriented in the opposite direction similar interactions were also present with many of the same residues, but the characteristic ARG143 guanidino group nitrogen atoms were instead interacting with other hydroxyl groups on the ligand. The nearby residue ARG82 was also found to make a strong appearance in the ligand interactions, though its guanidino group nitrogen atoms were found to hydrogen bond more with hydroxyl groups associated with the pyridine ring of these vitamin B6 metabolites. The polar residues SER185 and HIS368 also commonly appeared in the pyridine ring associated hydroxyl groups’ hydrogen bonding ligand interactions with a smaller number of the ligand poses.

**Figure 7:**
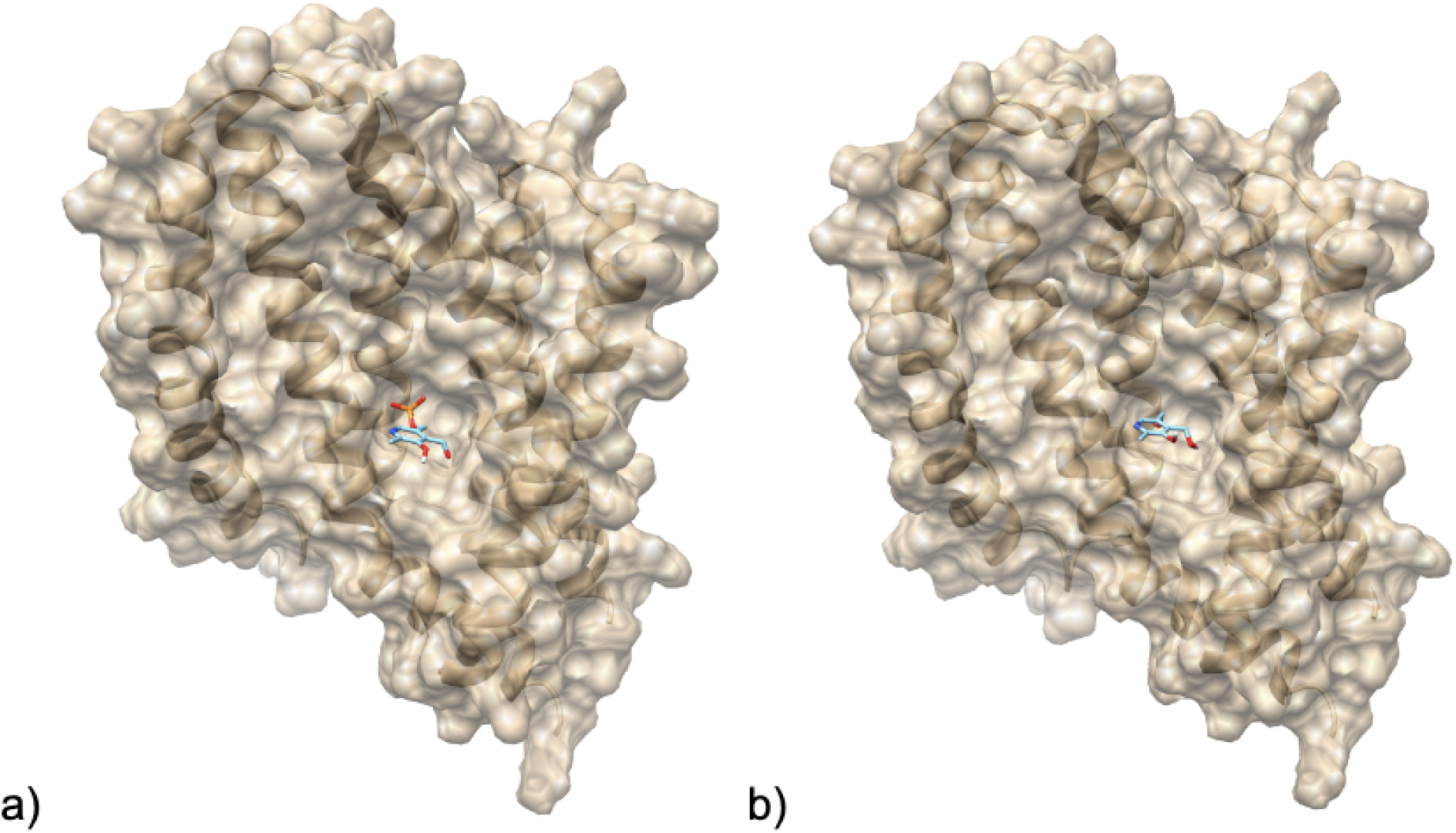
Docking ligand poses of Vitamin B6 to *Pf*_UVBSp. Depiction of the Webina second ranked PLP (a), and third ranked PN (b) poses in the pocket of the *Pf*_UVBSp homo-dimer top model.

Importantly, both ARG82 and ARG143 in *Pf*_UVBSp were found to be strongly conserved by CONSURF, though of course as seen previously in figure 5b, virtually all of the residues exposed at the interior surface are at least moderately conserved. We have anyhow additionally compared some of the other observed ligand pose-interacting residues to the equivalent residue positions in the Pfam seed MSA based on the alignment’s presence of RUS4 from *A. thaliana* [37]. ARG143 and HIS368 located at the back of the pocket also correspond to family-conserved positions of arginine and histidine respectively. However, ARG82 and SER185 located more towards the edge of the pocket were instead found to correspond to non-conserved positions where different types of residues can exist in the family. One unexplored idea based on this is that ARG143 and HIS368 may serve as residues required for general ligand binding by this domain, but residues equivalent to ARG82 and SER185 may instead provide ligand specificity determining sites within the family.

Ultimately, with regards to the RUS protein family as potential transporters of vitamin B6, AFDB models of the RUS proteins 1-6 from *A. thaliana* in-fact display variations in surface electrostatic charge [32], and do not all possess positively charged interior surfaces like *Pf*_UVBSp. One plausible explanation of this may arise from the fact that the three vitamers have a net charge of 0 (PL and PN) or +1 (PM) at physiological conditions, and that different members of the family simply transport different forms of the vitamin. Our docking results revealed the vitamin to be a highly feasible substrate for *Pf*_UVBSp, though of course even different proteins within the same family may transport different substrates altogether. Furthermore, it cannot be ruled out that the RUS proteins in *A. thaliana* are simply transporting a different metabolite responsible for the observed effects in plants occurring with some form of an interplay with the vitamin’s homeostasis. It must also be remembered here that in alignment with the positively charged pocket region, many of the other metabolites from the Kloehn *et al.* compiled list of apicoplast metabolites of unknown transporter are also negatively charged [23], and that the potential transport of another such metabolite cannot be ruled out also.

Lastly, it must be highlighted that the gene mutability index score reported by Zhang *et al.* for *Pf*_UVBSp in the asexual blood life stage was 1.0 (on scale from zero to one) [29], indicating that at this life stage the protein serves a much less critical transport function. In alignment with the previously discussed idea of the RUS family mediating homeostasis of the vitamin’s levels from serving as secondary transporters; *Pf*_UVBSp may similarly have more of a purpose in the apicoplast of *Plasmodium* species to maintain a metabolite’s levels.

### Pf_Memtrans

#### *Pf*_Memtrans domain structure

In alignment with the nine TMHs predicted by TOPCONS2 [84], *Pf*_Memtrans AF2 models adopted to a relatively large transmembrane domain structure of 10 TMHs (figure 8b) characterised by a few intricate structural features with regards to some of these individual TMHs. Most critically, as represented in figure 8c the core fourth and ninth TMHs of the domain crossover by a disrupted middle section lacking alpha-helical secondary structure (previously referred to as BHs). Also important in regards to potential homo-dimerisation, the first, second and seventh TMHs form the v-shaped scaffold region at the front of the domain. In terms of the surfaces of these AF2 models, two open funnel-shaped entrances lead into the transmembrane translocation region formed by the scaffold and core TMH-partitioning, which we here-onwards refer to as the tunnel region. We highlight here that most of the interior surface of this tunnel region is formed by the two BHs and TMH5 in the AF2 top model. The AF2 top model possessed the highest average pLDDT confidence score of 82.64, reflecting as seen in figure 8a that the transmembrane region was of high confidence and was the model used for all analyses.

**Figure 8:**
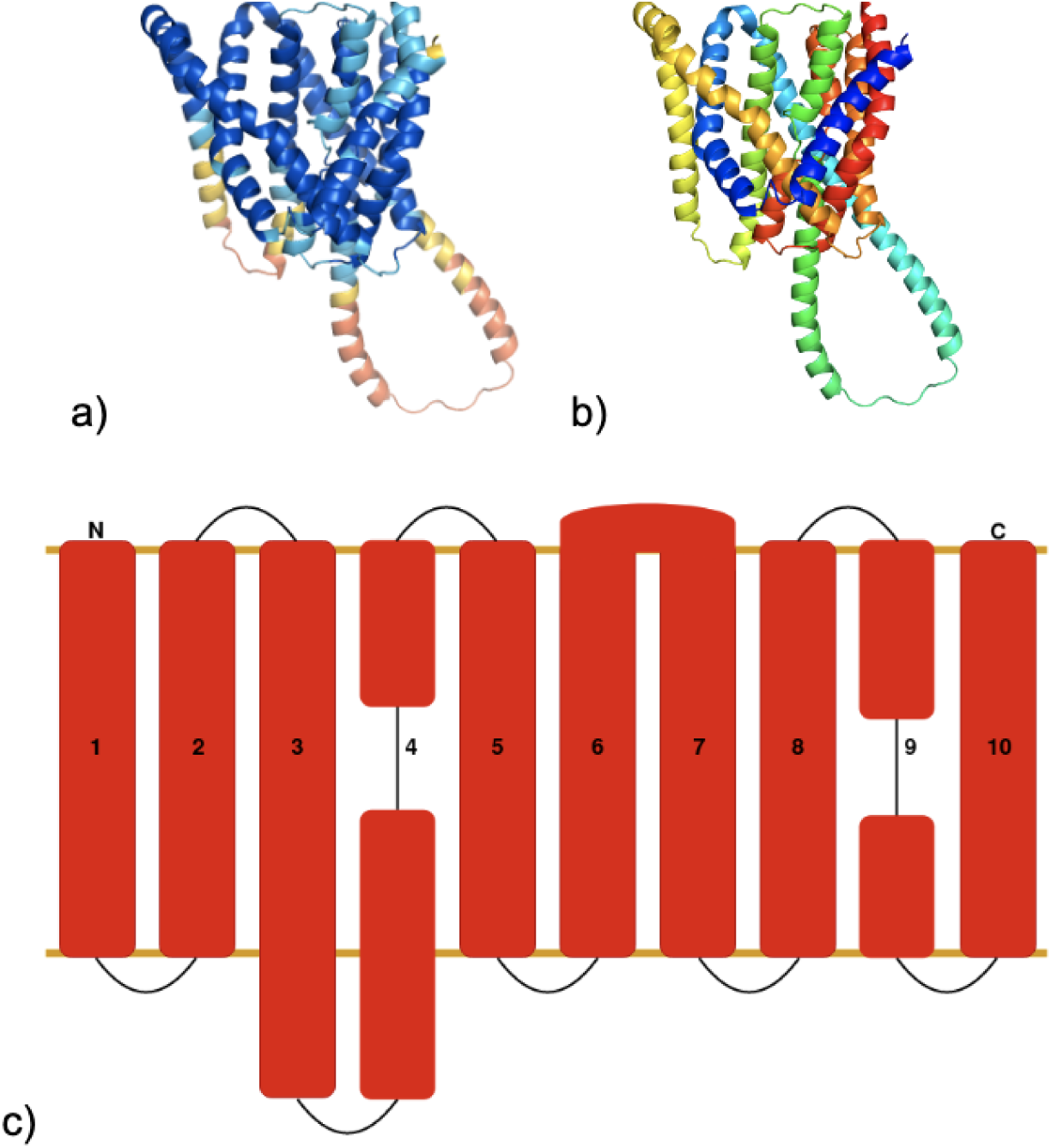
Predicted *Pf*_Memtrans structure and topology. *Pf*_Memtrans AF2 top model structure depictions: a) Model coloured by the previously described pLDDT confidence scheme. b) Model coloured from N-termini to C-termini (approximately colouring each of the TMHs 1-10 uniquely) by blue to red respectively. c) Cartoon *Pf*_Memtrans membrane topology by the same scheme as figure 1b.

DALI and GESAMT searches of *Pf*_Memtrans AF2 models against the PDBTM identified significantly similar CPAS structure hits, with the highest Z- and Q-score hits achieved being 20.7 and 0.25 respectively. The structure alignments in question covered virtually the whole of the hit and model chains of around 400 residues. All of the CPAS hits identified from these structure searches were members of the SBF, Na_H_antiport_1 and Na_H_Exchanger antiporter (Pfam) families. The use of TMalign revealed that these hits generally attained TMscores of close to 0.6, indicating that Pf_Memtrans possesses the same CPAS superfold. In line with this, at the time of conducting this research the model topology was found to be in perfect agreement with the Mem_trans domain topology proposed by Sudha *et al.* [61]. With the eventual release of the experimental structures of the AEC family [70–72], these models would also be revealed to possess uncannily high structural similarity to these structures down to the level of TMH bending as seen in figure 9a, plus would confirm the presence of the BHs. Here, a DALI alignment of the AF2 top model to the highest resolution structure 7wkw-A was marked by a Z-score of 18.8. Whilst we do not present the *Pf*_Memtrans AF2 homo-dimer models generated here, it is worthy of mentioning that: In alignment with the inter-monomer binding mode observed in the homo-dimeric AEC experimental structures (figure 9b), it was also observed that the monomers in a few of the homo-dimer models had also bound in variable ways around the same v-shaped scaffold region TMHs. Based on this and the other commonly observed homo-dimerisations of CPAS families by the scaffold region, it would appear that homo-dimerisation of *Pf*_Memtrans around this region as part of its activity is likely.

**Figure 9:**
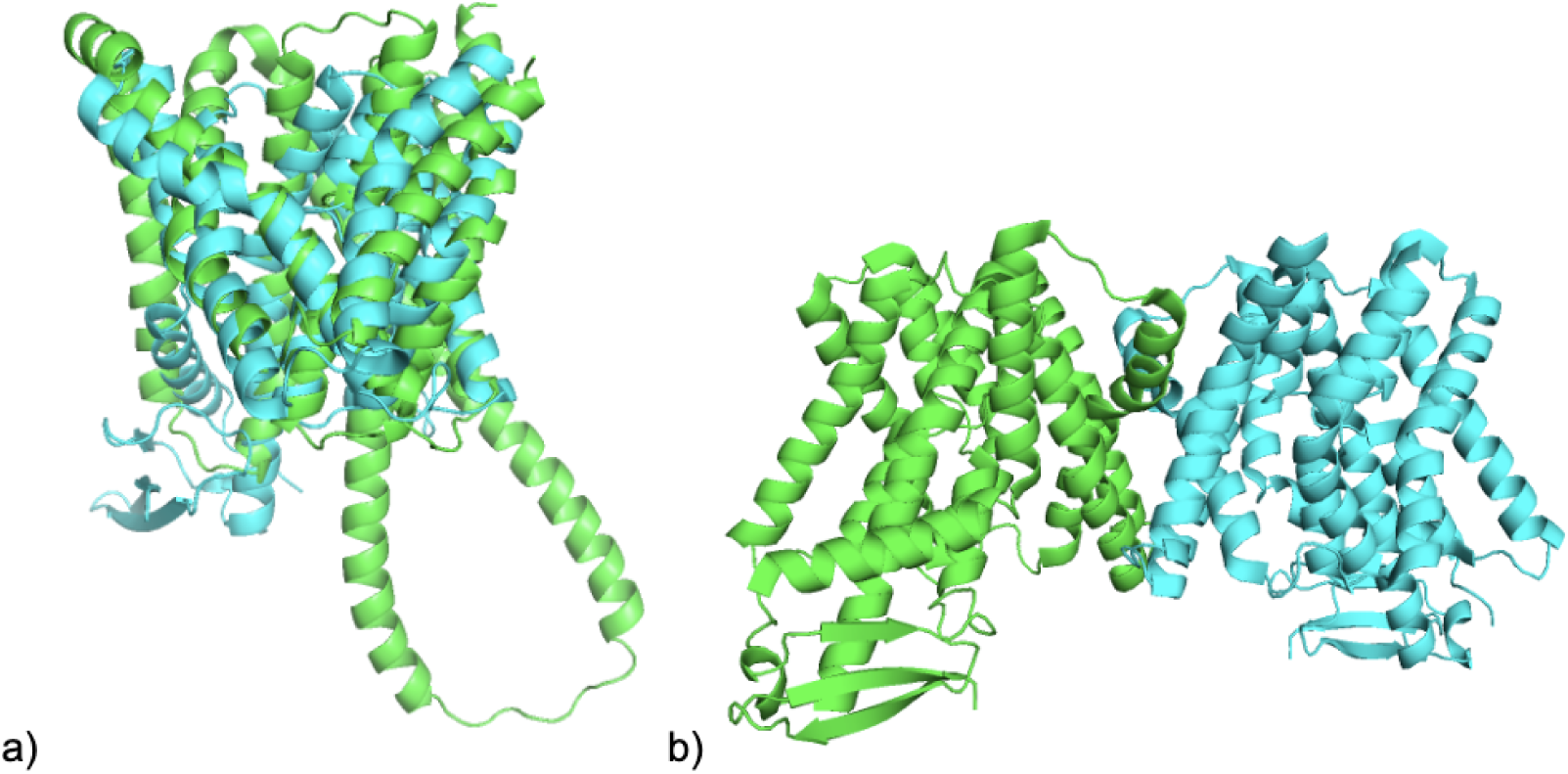
AEC family structure 7wkw. a) *Pf*_Memtrans AF2 top model (green) superimposition on PIN3 structure 7wkw-A (cyan) from DALI structure alignment [71]. b) 7wkw homo-dimer structure, chains A and B coloured green and cyan respectively.

#### *Pf*_Memtrans is a divergent member of the AEC family

Coinciding with the models’ theme of significant structural similarity to members of the CPAS, *Pf*_Memtrans was found to be homologous to a region of the Mem_trans domain representative PINI_ARATH (UniProt ID: Q9C6B8) identified by the HHsearch tool against Pfam, attaining a probability score of 80.74 and e-value of 0.1. Additionally, a small local region of similarity with the CPAS SBF domain was also identified here. The *Pf*_Memtrans sequence region of residues 310-460 that was aligned to PINI_ARATH corresponded approximately to the TMHs 6-10 (i.e., the C-terminal half) in the AF2 top model’s structure. Local regions of sequence similarity with CPAS members in the PDB in approximately the same C-terminal half region of *Pf*_Memtrans were not found by the HHsearch sequence search against the PDB70 database, but were by the equivalent AF2 run PDB70 search (used to identify structure templates). These corresponded to structures of the SBF, Na_H_Exchanger and Na_H_antiport_1 CPAS families.

Akin to *Pf*_UVBSp, though HHblits-UniProt and PSI-BLAST-NR sequence searches did not reveal the presence of homologues outside of *Plasmodium*, HHpred searches of the *A. thaliana* proteome identified three AEC family hits with e-values of less than 0.001 covering essentially all of both the query and hit sequence lengths. The highest similarity and scoring hit of these was the protein PIN-LIKES 7 (UniProt ID: Q9FKY4), with significant probability and e-value scores of 98.45 and 4.9e-04 respectively, though the alignment possessed a low number of identical residues as marked by a sequence identity of 13%. As the Mem_trans domain is the sole domain present in this family, based on these results *Pf*_Memtrans indeed possesses homologues in plant organisms. The other two HHpred proteome searches for this protein also revealed a lack of distant homology to other proteins which may contain the domain in *P. falciparum* and *T. gondii*, meaning that (as per *Pf*_UVBSp) within the Apicomplexa we have only identified single copies of the domain in *Plasmodium* species. We speculate and reason here that although there are currently no signs of the AEC family in chloroplasts, this protein’s ancestry would most likely be of cyanobacterium origin, based primarily on the presence of the AEC family in cyanobacteria [65]. However, with the somewhat ubiquitous nature of the family and domain, origins instead from the algal or secondary eukaryotic endosymbiont ancestors are also plausible, though further exploration here has been outside of our research scope.

#### *Pf*_Memtrans apicoplast metabolite docking screen

*Pf*_Memtrans was revealed from CONSURF analysis to possess a somewhat less strongly conserved transmembrane domain as reflected by figure 10a. However, at least a moderate or more proportion of the tunnel region’s surface lining is conserved as can be gauged from figure 10b, and is more positively charged as presented in figure 10c.

**Figure 10:**
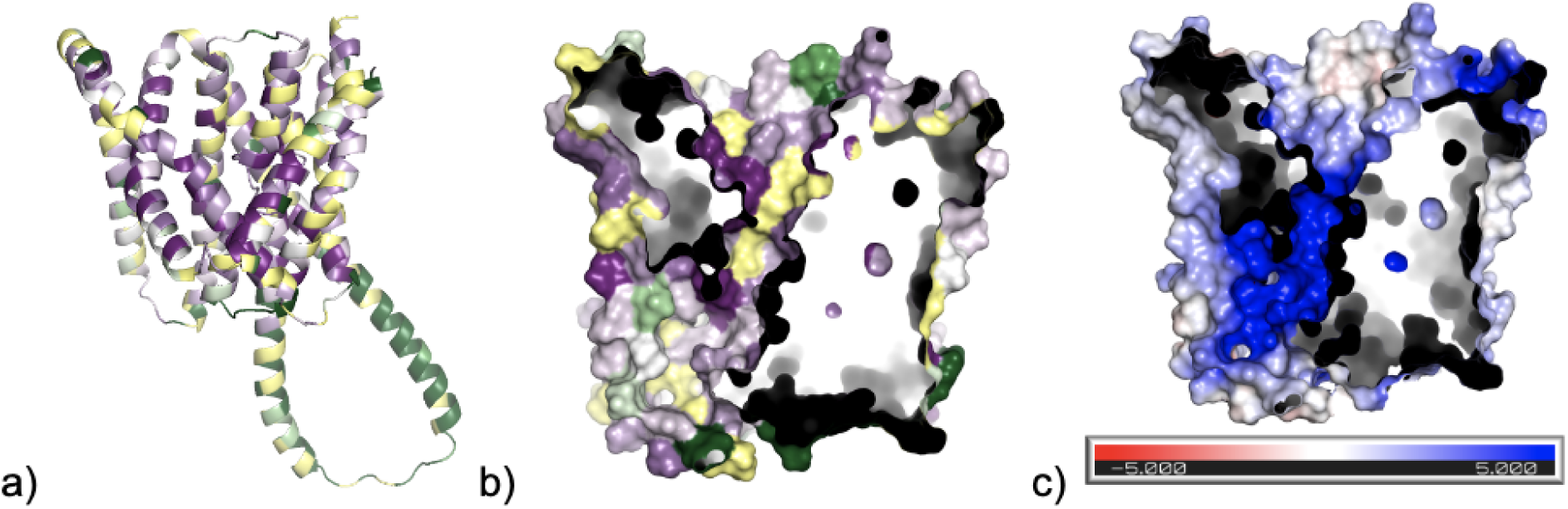
Conservation and internal surface electrostatics in *Pf*_Memtrans. AF2 top model coloured by the previously described CONSURF conservation scheme in a) cartoon form, and b) surface form with cross-section revealing the internal tunnel region. c) Surface electrostatics depiction of AF2 top model displaying similar cross-section.

For *Pf*_Memtrans we decided to dock a select set of the metabolites listed by Kloehn *et al.* (both imported and exported) currently lacking known apicoplast transporters to the AF2 top model [23]. Readers are encouraged here to consult the primary figure and its following caption in the Kloehn *et al.* publication in order to see this entire compiled list. This list of metabolites vary in size, molecule type and other chemical properties.

Importantly however, most of the metabolites have at least a slightly negative net charge. We firstly excluded the metabolites with net positive charges and the notably larger molecules including lipids and the cofactor Coproporphyrinogen III which were thought to be too large to be transported by *Pf*_Memtrans. Due to reasons of time we were not able to conduct docking to *Pf*_Memtrans of every metabolite belonging to a particular type of molecule in this list (namely ribonucleic acid and amino acid related), hence our *Pf*_Memtrans SMDS was not exhaustive. PLP was added as an additional candidate to the *Pf*_Memtrans SMDS due to the aforementioned evidence of its presence and lack of internal synthesis in the apicoplast [50,54]. The metabolites used in this SMDS (and their abbreviations) are all listed previously in table 2.

As displayed in figure 11, in the *Pf*_Memtrans SMDS it was observed in terms of the SWISSDOCK, CSM-lig and Prodigy-ligand PBAs of the tunnel region ligand poses that the larger and highest complexity metabolites DeP-CoA, CoA, NAD+ and NADP+ attained the best PBAs in comparison to the rest of the metabolites. In contrast, the five smaller metabolites of 5-ALA, ASP, GLU, aKG and citrate generally possessed the poorest ranges of PBAs from all four PBA sources out of the 16 metabolites docked.

**Figure 11:**
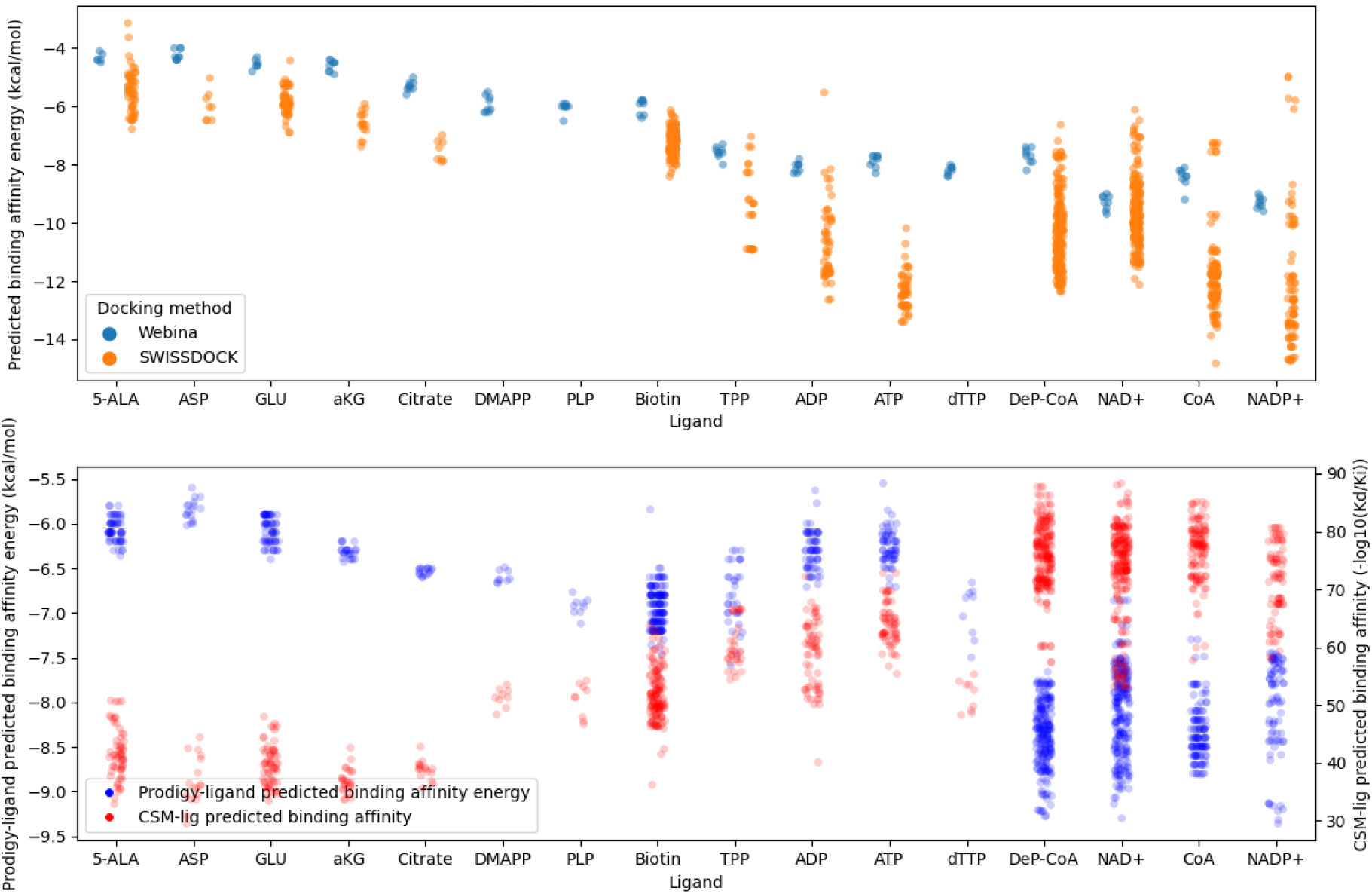
PBAs of *Pf*_Memtrans SMDS. Metabolites are ordered from left to right by increasing ligand complexity; all tunnel region poses from the *Pf*_Memtrans SMDS are depicted in both graphs. On the lower graph the very few ligand poses with negative CSM-lig PBA values have been omitted for both PBA types.

The remaining metabolites of DMAPP, PLP, Biotin, TPP, ADP, ATP and dTTP were more variable in the range of the PBAs their ligand poses attained. Though SWISSDOCK did not produce ligand poses of DMAPP, dTTP and PLP bound within the tunnel region, their corresponding Webina ligand poses also did not fair as well in the comparison of PBAs to the other larger and complex metabolites anyhow. Critically, it must be stated here that the PBAs attained in the tunnel region do not correlate with the ligands’ net charges, hence more negative net charges on the ligands are not thought to contribute to better PBAs in this region. The observation of the higher PBAs attained by the more complex ligands is likely due to the increased number of intermolecular interactions their ligand poses in the protein-ligand complexes possess. Though, in conjunction with this concept it cannot be ruled out that the native metabolite transported by this protein still possesses a higher molecular complexity; and conversely, the higher PBAs attained by these larger and more complex ligands may distract from the fact that a smaller and lower complexity metabolite with less prominent but still viable PBAs may be the native transport substrate.

We do not describe the ligand binding modes observed, except shortly in comparative terms to the observed binding modes of auxin (Indole-3-acetic acid, IAA) and inhibitor analogues in the AEC experimental structures. Ultimately, no particular trends in the PBAs of binding modes were ascertained, except for the following: The ligand poses which had bound lower in the tunnel region possessed markedly less negative Prodigy-ligand PBA energies. Furthermore, slightly more negative Prodigy-ligand PBA energies were attained by the ligand poses of the larger and more complex metabolites where their phosphate groups were positioned on-level with the residue LYS286 on TMH5. In conjunction with this, in the metabolite ligand poses where at least part of the ligand was positioned on-level with the grid centre residue (HIS247), the ligands had more interactions with residues on the BHs, and had a marginal tendency to attain some of the most negative Prodigy-ligand PBA energies observed. However, this happened essentially only for the larger and more complex ligands such as shown for the example ligand pose of DeP-CoA in figure 12b, and these higher affinity PBA energies are probably due to the ligand pose possessing more (favourable) inter-molecular contacts with residues. Critically, many of the residues are conserved on these BHs, and the conserved residues which commonly appeared in the ligand interactions as found by LigPlot are shown in figure 12a.

**Figure 12:**
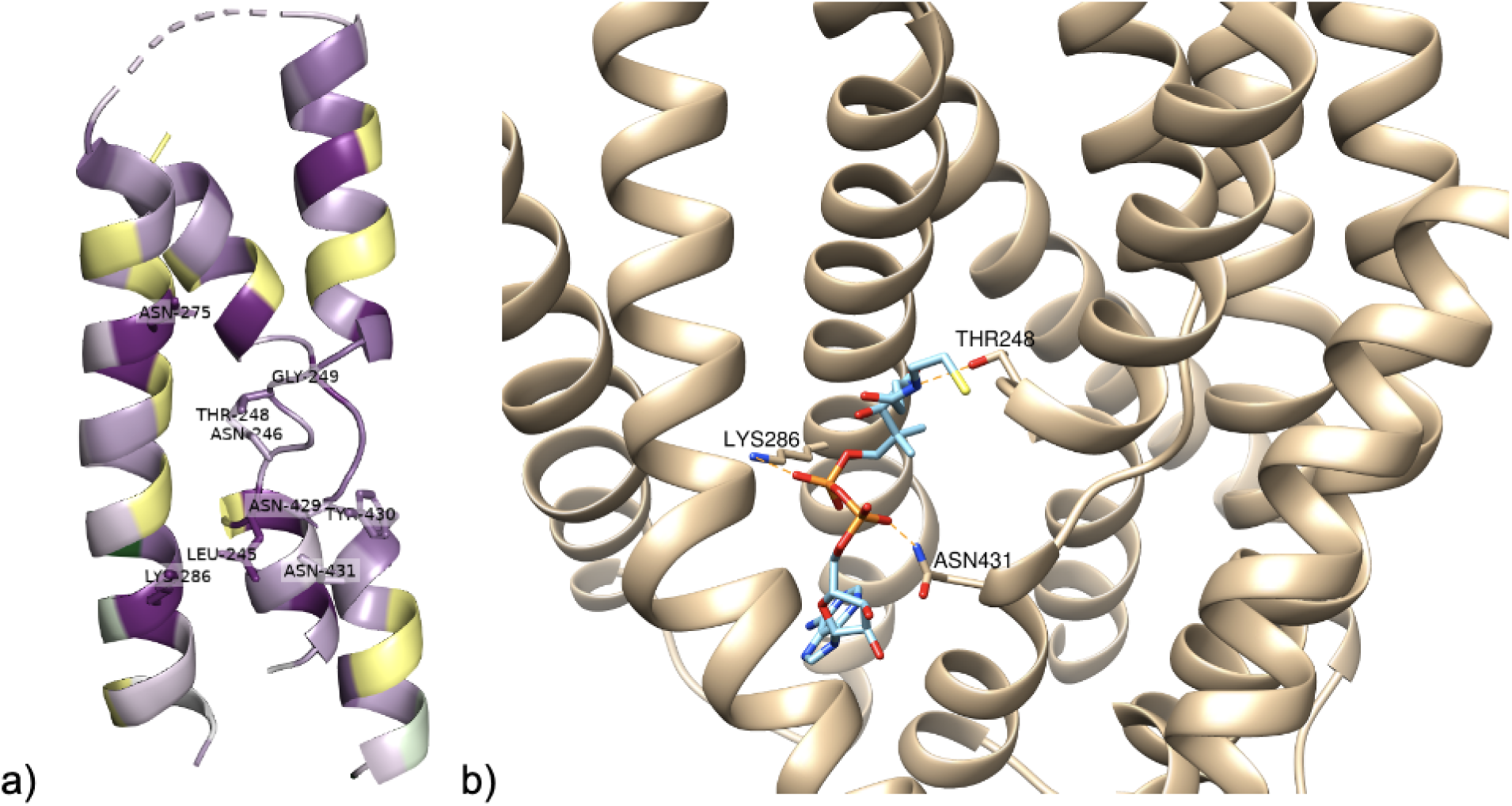
Likely ligand-interacting conserved residues of *Pf*_Memtrans tunnel region. (CONSURF-derived) Conserved residues found to be commonly involved in the LigPlot identified ligand interactions of poses from the SMDS. b) Structural depiction of the LigPlot identified residue-ligand interactions (marked by dashed orange lines) of the conserved tunnel residues (shown in a)) which appeared in the example of SWISSDOCK DeP-CoA top-ranked pose of cluster 55. Hydrogen atoms have been omitted for visual clarity.

A number of residue-ligand interactions are described from the AEC experimental structures including hydrophobic and hydrogen bonding interactions [71,72], plus indirect hydrogen bonding interactions through water molecules [70]. Some of these interactions also involve residues present on other TMHs besides the TMHs 4, 5 and 9 which were chiefly considered in the *Pf*_Memtrans tunnel. A general binding mode of IAA can be observed in these structures by the crossover of the BHs (not shown), and the other inhibitors also bound in similar modes and positioning. Though none of the Webina and SWISSDOCK metabolite ligand poses in *Pf*_Memtrans had bound significantly similarly, we now describe the key similarities and differences:

1. In regards to the most critical and consistently stated (direct) hydrogen bond interaction, the carboxylate group of IAA is oriented towards and hydrogen bonds with the AEC family-conserved residue ASN112 (ASN117 in PIN8) in the middle of the first BH. The structurally equivalent asparagine residue in *Pf*_Memtrans corresponds to ASN246, suggesting that this same residue (also conserved in *Pf*_Memtrans) may also enable an important hydrogen bonding interaction with the native substrate. In terms of relatable observations in the ligand poses, Webina-derived poses of mostly only the smaller metabolites with carboxylate groups where these CO2-groups were oriented towards this part of the first BH in a similar manner, though a similar occurrence for phosphate groups was less common.
2. The ligand carbon-ring groups were described to possess hydrophobic contacts involving a region of the tunnel involving at least two important hydrophobic residues corresponding typically to two consecutive valine residues just below the start of the broken segment of the second BH. In *Pf*_Memtrans, the structurally equivalent residues are instead polar and correspond to TYR430 and ASN431. Hydrogen-bonding interactions in the ligand poses of metabolites with ring-type groups were occasionally observed involving the ring groups and these two residues. However, these interactions mostly involved the ribose-ring groups present in the larger and more complex metabolites and rarely corresponded to other ring-moieties. Anyhow, one possibility here is that these two conserved polar residues in *Pf*_Memtrans may hydrogen bond with another type of polar chemical moiety on the native substrate (potentially on the some of the ring groups present in the metabolites), instead of the hydrophobic contacts observed in the AEC structures.
3. The residue TYR145 (GLN145 in PIN8) on TMH5 was implicated in stabilising ligand-residue hydrogen bonds and it is structurally equivalent to residue LYS286 in *Pf*_Memtrans. Interestingly, inspection of the Pfam seed MSA reveals that this sequence position (from structural equivalency in PINI_ARATH) appears to be a well-conserved tyrosine residue in the AEC family. One idea here based on the previously described interaction of ligand phosphate group interactions with the LYS286 side chain is that this conserved positively charged residue may instead act in this *Plasmodium* orthologue cluster to stabilise negatively charged ligand groups such as phosphates at this site.

Though such an open screening approach did not suggest a single transport substrate; based on the alignment of *ii*) and *iii*) together with the previously described higher PBAs plus the increased number of BH-residue interactions attained by ligand poses; we infer that *Pf*_Memtrans likely transports a larger and higher complexity metabolite with one or more phosphate groups. Importantly, it must be remembered that a number of the other metabolites listed by Kloehn *et al.* which were not screened also possess these chemical attributes (namely ribonucleic acid related). Other currently unknown imported or exported apicoplast metabolites may also be viable substrates in physico-chemical terms. Other clues to the transported metabolite may revolve around the fact that though *Pf*_Memtrans was also not found to be essential by Zhang *et al.* [29], it was found to have a low mutability index score of 0.27, close to that of 0.17 found for the essential UA6TM, and thus may transport a more critical apicoplast metabolite in the parasite’s blood life stage.

### UA6TM

#### Deep-learning structure predictions reveal a core six-TMH domain

Two structural regions were present in the UA6TM AF2 models (figure 13a) corresponding to a large alpha-helix of around 60 residues and a downstream transmembrane domain of six TMHs containing the sequence location of all four TMHs predicted by TOPCONS2. We refer to the former structural feature as the Large alpha-Helix Region (LHR) and abbreviate the latter domain as the C-terminal alpha-Helical Domain (CTHD). The distinction of these two structural regions is also validated from the fact that CONSURF analysis (figure 13b) had also revealed only the CTHD to be conserved in this protein. Also in these models, the fifth TMH of the CTHD possessed a particular disruption along its length. Interestingly, though the AF2 models possessed a low average pLDDT confidence, our obtained RosettaFold models of UA6TM (figure 13c) had also produced a similar type of structure characterised by the two structural regions [34]. Albeit, in the RosettaFold models the two regions were less separate, plus the CTHD was more compact and lacked the disrupted TMH. The RosettaFold model confidence score here of 0.53 was also relatively low, though in terms of the per-residue C-alpha estimated error (Å) seen in figure 13c, much of the CTHD was of lower than 5Å indicating that this region’s structure is of reasonable model quality.

**Figure 13:**
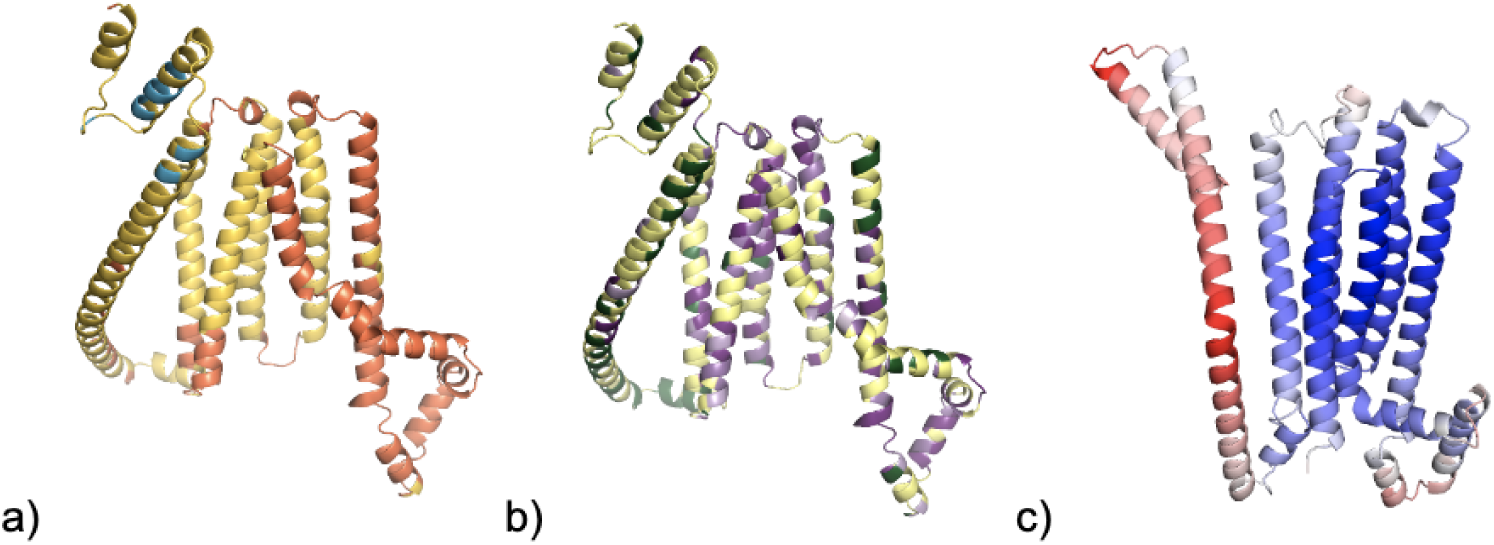
AF2 and RosettaFold models of UA6TM. UA6TM AF2 top model residues coloured by a) pLDDT confidence, and b) CONSURF conservation schemes. c) UA6TM RosettaFold top model coloured by blue to white to red spectrum for per-residue C-alpha estimated error in Å, corresponding to a lowest value of 1.75 (blue) to a highest value of 16.32 (red) respectively.

Although the AF2 models of UA6TM had attained some structure hits in the PDBTM with DALI Z-scores as high as 10.5; the superimpositions of these hit chains on the models displayed only partial similarity of a few alpha-helices to the CTHD, or corresponded to where the LHR had superficially been aligned to some longer TMHs in membrane protein structures. Due to this we had also conducted searches of these models with the LHR removed, but still arrived at the same result. Thus, no fold matches were identified from these models. The closest theme of a structure match identified from the main model structure searches was for the RosettaFold top model (with its LHR removed) with the cytochrome b561 family which also possesses six TMHs. Here a highest scoring hit found by GESAMT of Q-score 0.19 corresponded to 4o6y-A, a score which would normally indicate reasonable similarity. However, the structural similarity upon closer inspection of the superimposition was found to be questionable despite an approximately identical topology. Additional searches of the AFDB via the FoldSeek webserver of the AF2 top model also ultimately did not identify any hits with viable similarity to the CTHD [32,90]. As these structure searches were conducted a few years earlier in our research, shortly prior to the submission of this preprint we have thus conducted an additional up-to-date structure search of the entire PDB via the DALI webserver to check that no experimental structures with similarity have been released since (and was confirmed) [92].

The poorer confidence UA6TM AF2 models highlight that some struggles in next-generation structure prediction methods still exist. Although high accuracy at low MSA depth was reported for AF2 [1], the developers also reported a MSA depth of 30 sequences to still be a critical point likely necessary for accuracy, as covariance information is still utilised to predict information such as inter-residue distances. For UA6TM we have additionally conducted more recent structure predictions by use of the ColabFold AF2 webserver (see methods) [86]; a prediction which happened to be generated by an MSA with a greater number of 38 *Plasmodium* sequences (from 12 in the original AF2 run). However, the resulting AF2 model structures here were again highly similar to the previous AF2 models and of low pLDDT confidence. Such an observation highlights that other key factors here for UA6TM may involve the fact that: *i*) There may be a paucity of transmembrane protein structures in the PDB with similar native structural features that the AF2 deep-learning networks have been trained on. *ii*) The AF2 structure prediction process does not possess intelligence of molecular environment effects such as the membrane bi-layer and potentially unknown protein-protein interactions.

#### Proposed UA6TM membrane topology

Despite the lack of identified homologous structures the online PDBTM database was manually searched to visually identify analogous membrane protein structures possessing a six TMH domain and a separate large alpha-helix [78]. Only one conspicuous protein, “Ion channel TACAN” was identified corresponding to the structures 7n0k (figure 14a) and 7n0l [109], though the putative ion channel function was called into question by another study [110]. However, its long alpha-helix does not exist positioned vertically parallel to the transmembrane domain but lies horizontally below the membrane as it mediates homo-dimerisation (figure 14b). The AF2 Homo-dimer models of UA6TM presented highly similar (and very low pLDDT confidence) predicted monomer structures to the previously described AF2 models, binding mostly by TMHs of the CTHDs. However, interestingly in both the default and increased number of samples structure prediction runs, a small proportion of the models corresponding to lower ranked models instead possessed the monomer chain structures where the LHR was instead positioned below the CTHD in a horizontal manner. These bore some resemblance to the previously described example of the TACAN structure such as the prominent example displayed in figure 14c. Superficially, the other UA6TM models would resemble a transmembrane protein of seven TMHs. However, in regards to such a hypothetical alternative horizontal orientation of the LHR in the native structure, other observations from our research support the idea that no parts of the LHR are transmembrane spanning due primarily to a lack of TMH-prediction and hydrophobicity. We do not present or discuss this further due to the scope of this preprint, though this would suggest that in these models the LHR has been misoriented with respect to the CTHD. The previously described effects of the molecular environment in the above subsection plus potentially others may even explain this hypothesised mispositioning of the LHR in the prediction of this protein’s structure.

**Figure 14:**
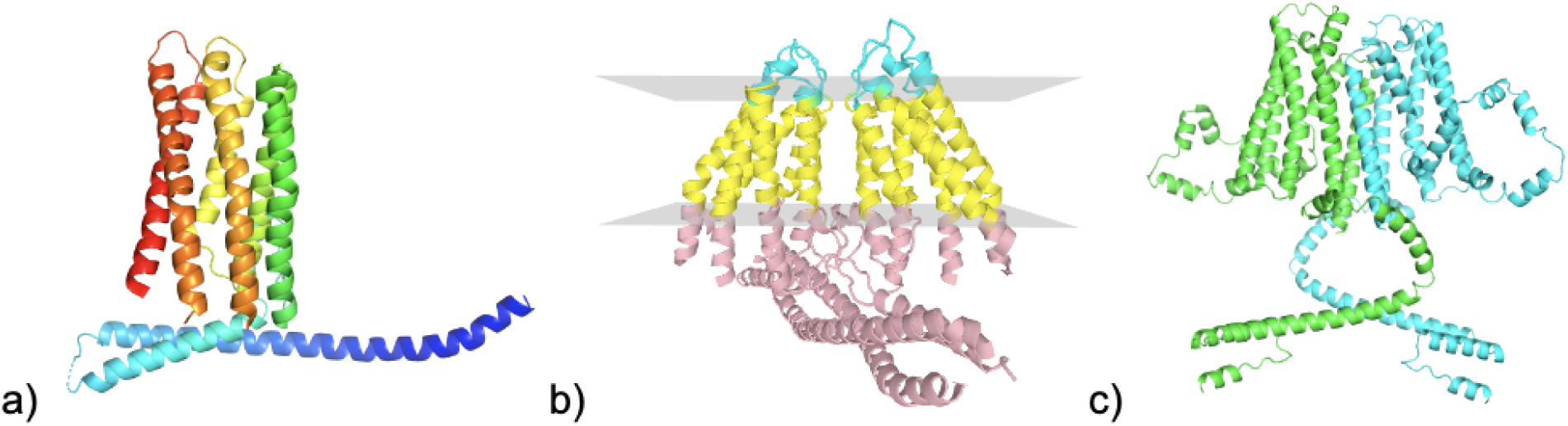
TACAN structure 7n0k and UA6TM alternative LHR orientation. 7n0k-A coloured from N-terminus to C-terminus by blue to red respectively. b) PDBTM membrane positioning prediction for 7n0k homo-dimer [78]. Planes of the membrane are grey, with extracellular, membrane and intracellular chain regions coloured cyan, yellow and pink respectively. c) The UA6TM 73rd ranked top model from the AF2 homo-dimer structure prediction where sampling was increased.

#### UA6TM as a hypothetical transporter

Unfortunately, from the previously described range of conducted bioinformatics sequence searches a lack of distant homology to known domains and PDB structures was found for this protein in addition to the lack of homologues outside of *Plasmodium*. This means that no subtle sequence-hints possibly pertaining to function can be further speculated on. However, whilst the CTHD region of UA6TM models lacked a transmembrane fold match, it could be posited that simply based on the presence of six TMHs approximately forming a simple alpha-helical bundle with a (appropriate sized) hollow, that this region could act as a transmembrane channel to transport an inorganic ion or small metabolite. The observed positive electrostatic charge in the interior surface lining between the TMHs in the AF2 models (not presented) would suggest negatively charged substrates if it did have a transporter function. Another similar possibility here is even to enable the transfer of electrons in a case similar to that of the previously described cytochrome b561 structure hits as redox activity has been confirmed in the apicoplast [111]. Due to the apparently essential nature of this protein in the blood life stage [24,29], the molecular function may enable an important niche function of the apicoplast present in these members of the *Plasmodium* genus that may not be present in other members of the Apicomplexa.

## Conclusions

Due to the previously discussed biological plant protein contexts of the domains present in *Pf*_UVBSp and *Pf*_Memtrans respectively, we propose there is a likely chance that these genes have originated from ancestor proteins in the same cyanobacterial membrane(s) in the evolutionary history of the Apicomplexa. Such ancestry would now expectedly place the specific membrane-residency of these transporters at least within the two innermost apicoplast membranes, though how transporters operate the required metabolome transport through all four membranes is poorly understood. The lack of the appearance of homologues in other Apicomplexan species outside of *Plasmodium* may anyhow suggest that during the evolution of the Apicomplexa after the final symbiosis event that either: *i*) Despite the reduction of the ancestral genes, these transporters have been kept in the *Plasmodium* lineage from either a maintenance or adaptation of an original transportive function. Or *ii*), a strong divergence of these transporter families occurred within the phylum to the point of a lack of recognisability. What both of these ancestor transporters were originally transporting is not known and the extent to which the sequence differences between the *Plasmodium* and plant protein families affects the similarity of the native substrates transported is unclear.

The presented *in silico* discovery of *Pf*_UVBSp and *Pf*_Memtrans as proteins with a transporter molecular function belonging to known superfamilies will provide a bridge for apicoplast bioscientists to experimentally characterise them further, ultimately helping to better understand the apicoplast. Additionally of importance, the knowledge of the UVB_Sens_prot domain as a MFS transporter forming component will secondarily enable plant molecular biologists towards a better understanding of RUS proteins in vitamin B6 homeostasis. Similar future investigations are now warranted for other uncharacterised apicoplast transmembrane proteins, which may again successfully reveal the presence of other transporter domains present in plant protein families due to the apicoplast’s ancestry. With regards to the lower confidence but reasonable transmembrane structure prediction characteristics of UA6TM, though AF2 has been documented with vast arrays of successful examples of protein structure prediction, the case of some proteins such as this exemplify its limitations. However, the further development and application of such predictive structural bioinformatics methods like AF2 will be instrumental in illuminating other critical proteins in disease-causing microorganisms.

